# A versatile, positive-going voltage indicator that enables accessible two-photon recordings in vivo

**DOI:** 10.64898/2026.04.07.717088

**Authors:** Alex James McDonald, Michelle Ann Land, Shuyuan Yang, Noura Hakam, Vincent Villette, Jun Zhu, Mario Galdamez, Mario Fernandez de La Puebla, Xiaoyu Lu, Gregory Foran, Tanmayee Torne-Srivastava, Beatriz Campillo, Haixin Liu, Xiaoyu Dong, Shujuan Lai, Matthew Shorey, Hala Abdallah, Raegan Banks, Anastasia Mamontova, Yu-Yau York Shan, Ryan Kroeger, Robert G. Law, Ming Hu, Daniela Gaspar Santos, Jonathan Bradley, Alberto Lombardini, Benjamin Mathieu, Annick Ayon, Ryan Gregory Natan, Hongyi Yuan, Jacob Reimer, Laurent Bourdieu, Na Ji, Weijian Zong, François St-Pierre

**Author notes:** Contributed equally to this work (Alex James McDonald, Michelle Ann Land, Shuyuan Yang). Contributed equally to this work (Noura Hakam, Vincent Villette, Jun Zhu, Mario Galdamez, Mario Fernandez de La Puebla, Xiaoyu Lu). To whom correspondence should be addressed: reagents C in-vitro data (FSP); resonant-scanning data (JR); ULoVE C diagonal scanning data (LB); FACED data (NJ); MINI2P data (WJ).

## Abstract

Genetically encoded voltage indicators (GEVIs) enable cell-type–specific optical readout of membrane potential, but two-photon (2P) spike detection has been hampered by low signal-to-noise and ultrafast off-kinetics, restricting use to specialized microscopes. We introduce FORCE1s, a green, positive-going GEVI engineered to make robust 2P voltage imaging broadly accessible. FORCE1s brightens from a dark baseline during depolarization, reports spikes with ∼100% ΔF/F in awake mice, and displays repolarization kinetics that are tuned for reliable spike detection at sub-kilohertz frame rates. As a result, FORCE1s supports spike-resolved multi-cell recordings on standard resonant-scanning microscopes, and further scales to larger fields of view and neuron counts on advanced modalities. FORCE1s also enables multiplexed voltage–neurotransmitter imaging and extended recordings in freely moving mice using a compact, affordable MEMS-based 2P miniscope. Together, these advances establish FORCE1s as a community-ready tool that democratizes deep-tissue voltage imaging across platforms and experimental contexts.

## Introduction

Membrane potential dynamics underlie neuronal computation, shaping how dendrites integrate inputs and how neurons generate action potentials that drive communication across neural circuits. Genetically encoded voltage indicators (GEVIs) provide a powerful optical approach for measuring these rapid electrical events with cell-type specificity^1,2^. Recent work has shown that two-photon (2P) optical recording approaches can capture spikes and subthreshold events across multiple neurons, but require specialized microscopes capable of ultrahigh sampling rates—systems available to a small fraction of neuroscience laboratories^3–8^. As a result, deep-tissue, multi-cell 2P voltage imaging remains inaccessible to most of the community, limiting both the reach and the scientific impact of GEVI technology. Overcoming this barrier requires indicators that perform robustly on widely used resonant-scanning microscopes and that also elevate the capabilities of advanced optical recording methods.

A central barrier to the broad adoption of 2P voltage imaging is the ultrafast repolarization kinetics of existing GEVIs. These kinetics force indicators like JEDI-2P and JEDI3sub/hyp to be sampled at near-kilohertz frame rates to detect action potentials with high fidelity^3,5,9^. Conventional resonant-scanning microscopes cannot routinely achieve these recording speeds within fields of view large enough for multi-cell recordings^5^, preventing meaningful co-modulation or circuit analyses. Even cutting-edge microscopy platforms struggle to scale with such rapidly responding indicators, inevitably imposing a trade-off between field of view (or cell count) and spike-detection accuracy.

A second major obstacle is the small fluorescence responses of current indicators. This limitation amplifies the demands imposed by fast kinetics: high sampling rates reduce photon counts per timepoint, leading to noisier baselines and undermining spike detectability. Neuropil background fluorescence further degrades signal quality^3^, often forcing even sophisticated imaging systems to rely on sparse expression strategies^5^.

Here we introduce FORCE1s, a green, positive-going GEVI engineered to address these two challenges. It achieves larger spike- and subthreshold-evoked responses while also offering high intrinsic brightness and improved photostability. FORCE1s’ repolarization kinetics strike a crucial balance: fast enough to resolve individual spikes, yet slow enough to support the subkilohertz sampling rates. These enhanced properties enable reliable, scalable voltage imaging on conventional resonant scanning microscopes while further enhancing the performance of advanced modalities, including ULoVE^4^, Diagonal Scanning, and FACED^8,10^. FORCE1s also enables 2P voltage imaging with miniscopes comparable to commercially available systems, empowering the broader community to conduct deep-tissue voltage imaging on freely moving animals. By supporting spike-resolved, multi-cell voltage imaging across both standard and state-of-the-art platforms, FORCE1s democratizes access to deep-tissue voltage imaging, opening the way for the broader neuroscience community to map voltage dynamics across larger populations and in more naturalistic behavioral contexts.

## Results

### Two-photon directed evolution of a positive-going voltage indicator

A promising strategy to increase spike detectability is to invert the response polarity, creating positive-going indicators that brighten during depolarization. For a given absolute fluorescence change, polarity inversion increases the fractional response amplitude (ΔF/F)^11^ and reduces baseline photon shot noise from both the neuron and the surrounding neuropil^12^, thereby improving the overall signal-to-noise ratio (SNR). Positive-going indicators can also exhibit greater photostability when their brightness modulation arises from changes in chromophore excitation.

We selected the bright-to-dim indicator JEDI-2P as the parental sensor due to its proven performance in multiple 2P voltage optical recording paradigms^5,7,8^ and the absence of other dim-to-bright indicators^13–16^ at the project’s inception. The high-throughput 2P HEK cell expression-based screening platform was then used to simultaneously assess response amplitude at sub- and suprathreshold voltages, baseline brightness, kinetics, and photostability^5,17^. In total, 21 saturation mutagenesis libraries targeting 12 residues within the voltage-sensing domain (VSD) and GFP were screened (Figs. 1A, S1.1).

**Figure 1.**
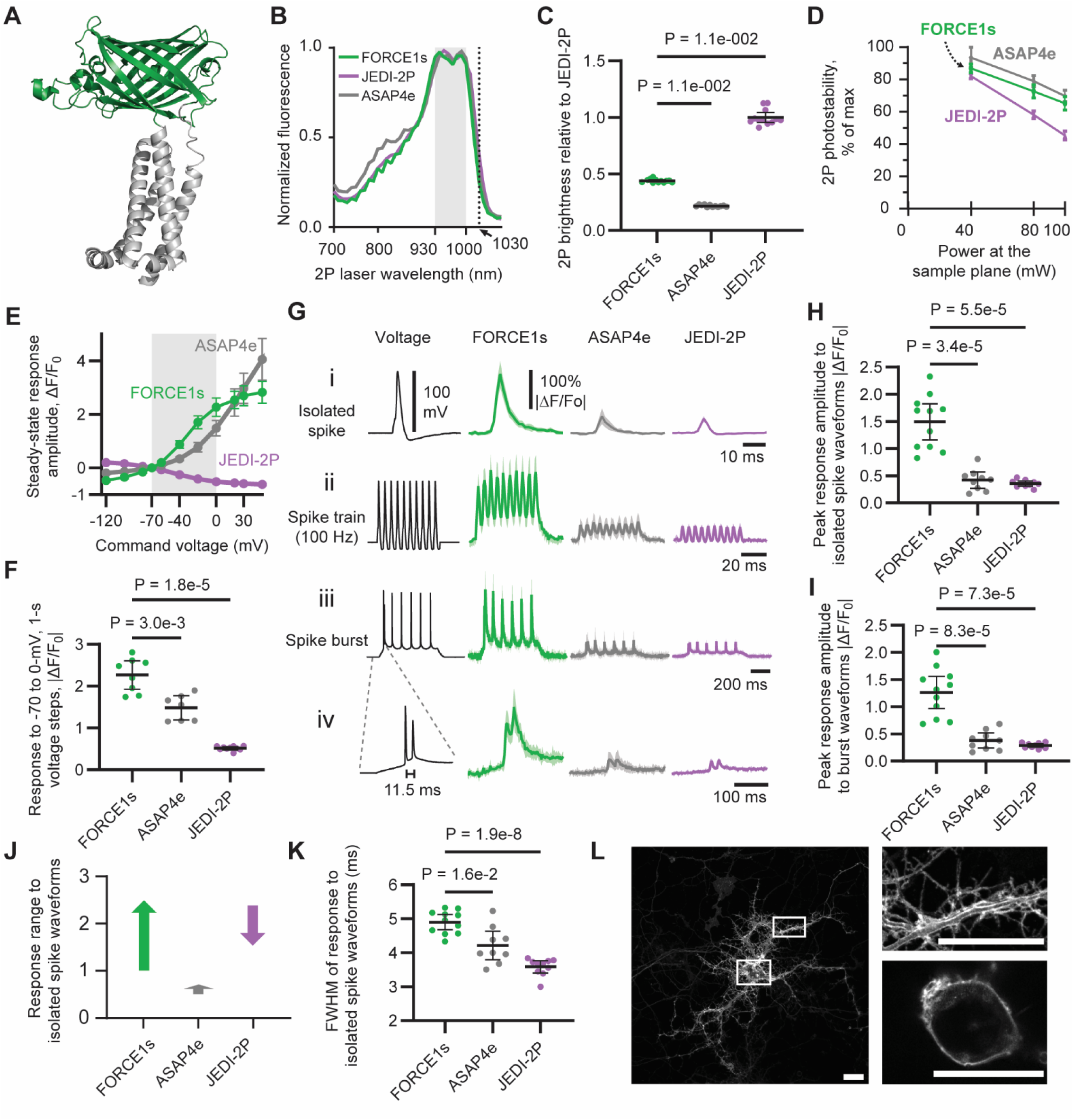
FORCE1s is a brighter, more sensitive 2P-optimized indicator tuned to physiological voltage ranges. All data were collected in mammalian cells under 2P excitation, except for panel L, which was recorded in 1P. (**A**) An AlphaFold3-predicted structural model of FORCE1s. The circularly permutated GFP (cpGFP) is colored in green and the voltage sensitive domain plus linkers are colored in gray. (**B**) 2-photon excitation spectra. Emission: 525/50 nm. Cell type: HEK293A cells. The gray-shaded area (930-1000 nm) spans the portion of FORCE1s’ spectra with >90% of peak absorption. n = 15 (FORCE1s), 16 (JEDI-2P), 15 (SpikeyGi2), or 12 (ASAP4e) fields of view of >100 cells. (**C**) Brightness comparison at 940 nm, −70 mV, and 22°C. GEVI and mBeRFP emission signals were ratioed to normalize for expression variation. Datapoints are the mean from n = 12 independently transfected wells/GEVI. Error bars: 95% CI. Black lines: per-GEVI mean. P-values from a Kruskal-Wallis test. (**D**) Photostability comparison at 940 nm, −70 mV, and 22°C. Values represent the area under the curve (AUC) of 6-min fluorescence-vs-time traces, expressed as a percentage relative to a non-bleaching GEVI. Datapoints show means from n = 12 independent transfections, each with >50 measured cells. Error bars: 95% CI. (**E-I**) Characterization via whole-cell voltage clamp and 2P imaging at 33 °C. (**E**) Voltage response curves, shown as the mean steady-state fluorescence response to voltage steps from −70 mV. The gray shaded area represents our target voltage range, from −70 mV to 0 mV. n = 7 (FORCE1s), 5 (ASAP4e), and 10 (JEDI-2P) HEK293A cells. (**F**) Quantification of panel E for 70-mV voltage steps. Black lines: mean. Error bars: 95% CI. P-value from Dunnett’s T3. (**G**) Responses to spike waveforms: (i) isolated spikes with 100-mV amplitude and 2-ms full-width at half maximum (FWHM); (ii) a 100-Hz train of i’s waveform; (iii) spike burst on a subthreshold depolarization. Spike width, 1.5-2-ms FWHM; (iv) zoom-in from iii. Imaging by resonant-scanning at 721 Hz. Dark traces: mean. Shaded areas: 95% CI. n = 10 (JEDI-2P), 9 (ASAP4e), and 11 (FORCE1s) HEK293A cells. (**H**-**I**) Peak response amplitude to isolated spikes (H) and spikes in bursts (I), quantified from G rows i and iii, respectively. Black lines: mean. Error bars: 95% CI. P-values from Dunnett’s T3. (**J**) Depiction of GEVIs’ change in brightness in response to the spike waveform from panel G (row i), relative to FORCE1s’ baseline. (**K**) Width of isolated spikes from panel G, row i. Black lines: mean. Error bars: 95% CI. P-value from Dunnett’s T3. (**L**) Representative confocal image showing FORCE1s expression and plasma membrane localization in a dissociated rat cortical neuron. The soma zoom-in was taken from a single confocal Z slice, while the other images are confocal Z-stack maximum-intensity projections. Scale bar: 20 µm.

We hypothesized that response polarity could be inverted by altering residues near the chromophore, thereby modifying how voltage-induced conformational changes modulate its brightness. Saturation mutagenesis of S151 —adjacent to the chromophore-interacting residue E152, a key mutation incorporated during JEDI-2P evolution— revealed that substitution with aspartic acid or proline inverted the response polarity. S151D yielded greater response amplitudes, while S151P showed higher baseline brightness. Given our primary objective of maximizing voltage response magnitude, we selected S151D for further optimization. In an example of convergent engineering, 151D was independently identified in ASAP4e^13^ and SpikeyGi^14^.

This final variant—harboring eight mutations relative to JEDI-2P—exhibited a 6-fold enhancement in response amplitude to long field stimulations, a 1.2-fold increase in peak brightness, and a 1.5-fold improvement in photostability in the context of screening assays. We designated this promising positive-going indicator Fluorescent Observer of Rapidly Climbing Electropotentials 1 (FORCE1s), with the ‘s’ designating both its greater sensitivity and its optimization for slower, larger-area voltage imaging.

#### FORCE1s is the brightest and most responsive positive-going voltage indicator

We benchmarked FORCE1s in vitro under 2P illumination (unless noted otherwise), comparing it to JEDI-2P and ASAP4e—the most responsive published positive-going 2P indicator^13^. We first determined FORCE1s’ 2P excitation spectrum to identify a range of wavelengths for efficient excitation. FORCE1s showed a broad excitation profile, achieving 90% of peak excitation between 930 and 1000 nm, with comparable peaks at 940 nm and 990 nm (Fig. 1B). Excitation at 1030 nm illumination was 26% of peak, enabling FORCE1s to be excited by powerful fiber lasers operating at or around this wavelength, as illustrated below with FACED microscopy. The excitation and emission spectra of FORCE1s, JEDI-2P, and ASAP4e were qualitatively similar (Fig. S1.2A,1B).

High brightness and photostability are essential properties for voltage indicators, ensuring reliable signal detection and compatibility with extended imaging sessions. In mammalian cells, FORCE1s was more than 2-fold brighter than ASAP4e (Fig. 1C). Although FORCE1s is a dim-to-bright sensor, it retained 39% of the brightness of the bright-to-dim indicator JEDI-2P (Fig. 1C). FORCE1s demonstrated enhanced photostability relative to JEDI-2P (See Suppl. stats). Despite its significantly higher brightness than ASAP4e, FORCE1s showed comparable or only slightly reduced photostability (Fig. 1D, S1.2B). The excited-state fluorescence lifetimes of FORCE1s and JEDI-2P showed negligible changes between depolarizing and hyperpolarizing conditions, suggesting that voltage primarily influences fluorophore excitation rather than emission (Table S1.3).

We next quantified depolarization kinetics (“on-kinetics”) of FORCE1s by measuring responses to voltage steps from –70 to +30 mV at 33°C (Table S1.4). These experiments were performed under 1P excitation, as 2P methods for accurately quantifying kinetics with sampling rates ≥10 kHz have not yet been established. ASAP4e’s depolarization kinetics were primarily governed by an 11-ms component, accounting for 66% of its response, while a faster 2.6-ms component contributed the remaining (34%). In contrast, FORCE1s exhibited a dominant 2.8-ms component, accounting for 93% of its response, enabling more effective tracking of brief voltage transients, such as action potentials. JEDI-2P displayed even faster on-kinetics, with a time constant of 0.49-ms, outperforming both positive-going indicators. Notably, all three GEVIs are expected to exhibit accelerated kinetics in vivo at 37 °C compared to the 33 °C conditions used in these assays, due to the temperature-dependent nature of their response dynamics^5^.

We characterized repolarization kinetics (“off-kinetics”) in response to steps from +30 to –70 mV. FORCE1s exhibited two time constants, 2.4 and 9.1-ms, each contributing approximately equally to the overall response. Compared with the 1.5-ms time constant of JEDI-2P, the slower off-kinetics of FORCE1s extends the photon collection window, enhancing spike detectability at a given sampling rate or allowing the recording of more cells. The 2.4-ms component of FORCE1s remains fast enough to support reliable action potential detection, even in spike bursts, as demonstrated below in vitro and in vivo.

We quantified the voltage response curves of GEVIs by measuring their steady-state fluorescence in response to voltage steps applied via whole-cell patch clamp from a resting potential of −70 mV. FORCE1s produced the largest responses across both subthreshold and spike ranges. For instance, its response to a 30-mV depolarization from −70 to −40 mV was 87.4 ± 13.9%, compared with 33.6 ± 6.4% for ASAP4e and 25.9 ± 3.7% for JEDI-2P (Fig. 1E, Fig. S1.2C). In the range of –70 to 0 mV, FORCE1s again showed the strongest response (227 ± 34%), outperforming ASAP4e (148 ± 29%) and JEDI-2P (52 ± 4%) (Fig. 1E-F). We focused on responses to 0 mV because, although action potentials can reach +30 mV or higher, the limited kinetics of GEVIs and the temporal resolution of typical microscopy setups reduce the contribution of voltages above 0 mV to the overall fluorescence signal.

We assessed each GEVI’s ability to report spike waveforms, which depend on both indicator kinetics and voltage responsivity. FORCE1s exhibited 3.6- and 4.2-fold greater responses than ASAP4e and JEDI-2P, respectively (Fig. 1Gi,H). Individual spikes were clearly resolved within 100-Hz trains (Fig. 1Gii) or superimposed on an extended subthreshold depolarization (“UP state”; Fig. 1Giii-1Giv). Consistent with their voltage response profiles, FORCE1s showed larger responses to the UP-state component of the waveform than both ASAP4e and JEDI-2P.

FORCE1s exhibited a larger dynamic fluorescence range in response to isolated spike waveforms—reflecting a greater change in brightness from baseline to peak—than both ASAP4e and JEDI-2P (Fig. 1J). Consistent with its observed slower off-kinetics, its response to spike waveforms was wider than ASAP4e and JEDI-2P (Fig. 1K). Finally, before conducting in vivo experiments, we confirmed that FORCE1s was expressed and trafficked to the plasma membrane of dissociated rat hippocampal neurons (Fig. 1L).

### FORCE1s-Kv detects spikes and subthresholds with high detectability in vivo

We began in vivo two-photon benchmarking of FORCE1s via simultaneous GEVI imaging and 2P-guided whole-cell patching in the visual cortex of anesthetized mice (Fig. 2A-B). FORCE1s was preferentially localized to the soma of neurons through the use of a Kv2.1-derived targeting tag, henceforth indicated by the suffix ‘-Kv’ in its name (Fig. 2B). Imaging at 440 Hz with a resonant scanning microscope, FORCE1s-Kv fluorescence closely tracked subthreshold voltage changes and reliably detected spikes, even during bursts (Fig. 2C-G). Notably, high spike detection fidelity was maintained across all cells using a single set of adaptive thresholding parameters, without per-recording adjustments (Figs. S2.1A-B). Using juxtacellular patch recordings, we found FORCE1s-Kv detected spikes with substantially higher fidelity than JEDI-2P-Kv, with mean F_1_ scores of 0.87 and 0.48, respectively (Figs. 2H-I).

**Figure 2.**
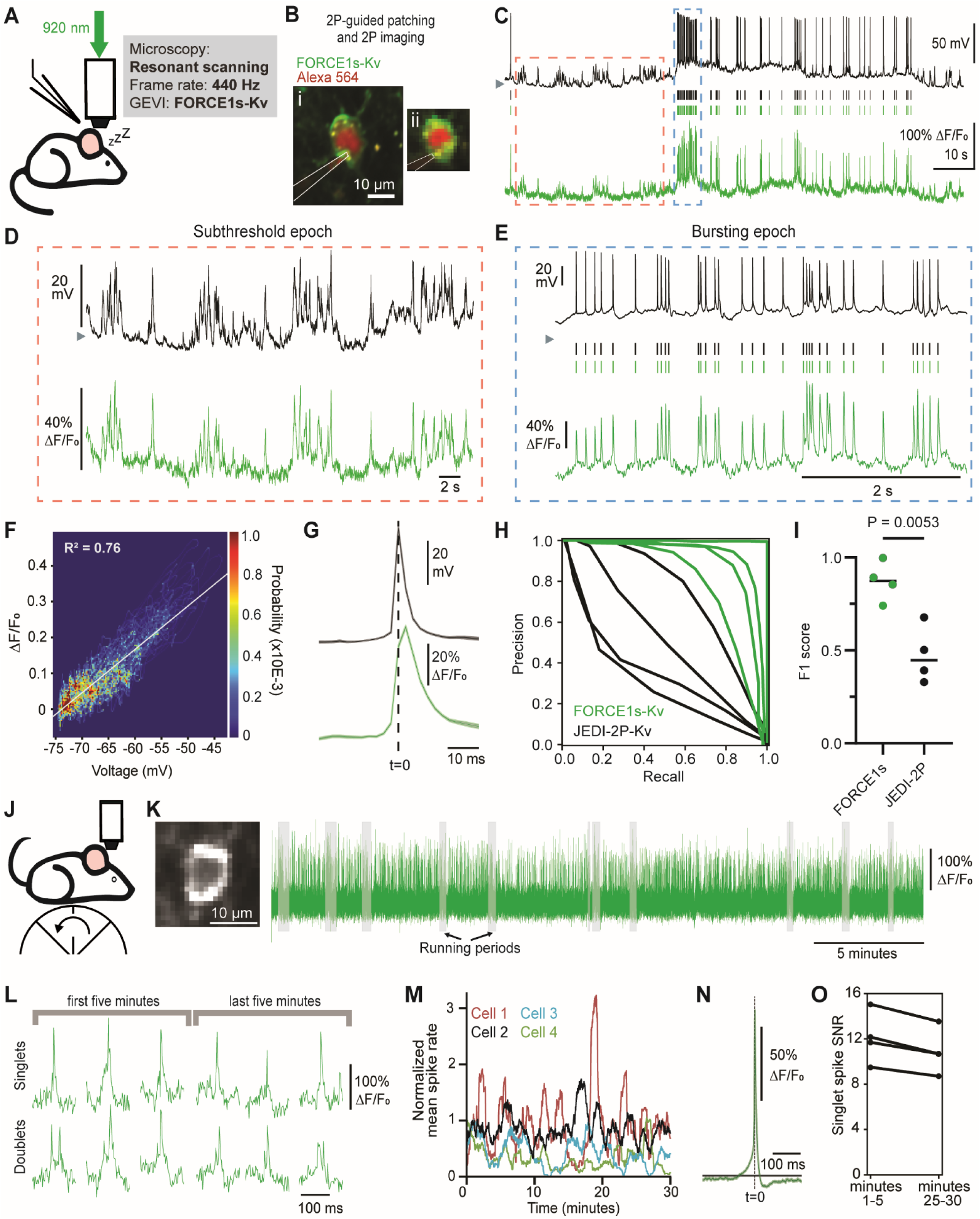
FORCE1s reports in vivo neuronal activity with high fidelity under resonance scanning 2-photon microscopy. (**A-I**) 2P-guided whole-cell patching and resonant-scan imaging in anesthetized mice. (**A**) Experimental schematic. (**B**) 2P images of a representative FORCE1s-Kv-expressing neuron are shown at high resolution (i) and the size and resolution used during voltage imaging (ii). The patching pipette is outlined. (**C**) Simultaneous electrophysiological recording (*top*, 10 kHz) and FORCE1s-Kv fluorescence imaging (*bottom*, 440 Hz). The fluorescence trace was low-pass filtered at 150 Hz to enhance the visualization of the spikes. Dashed boxes: epochs shown in panels D-E. Vertical lines: detected spikes. Gray triangle (*left*, and in panels D-E): −70 mV. (**D-E**) Zoomed-in subthreshold (D) and bursting (E) epochs. Panel D’s optical trace was low-pass filtered at 50 Hz to enhance the visualization of subthreshold fluctuations. (**F**) Probability distribution plot from panel D. Subthreshold sensitivity (linear fit): 0.012 ΔF/F_0_ per mV. (**G**) Electrical (*top*) and optical (*bottom*) spike-triggered average. n = 345 isolated spikes from the neuron recorded in panels B-E. Shaded region: SEM. (**H**) Precision-recall curves of juxtacellular patched neurons expressing FORCE1s-Kv (green) or JEDI-2P-Kv (black). The JEDI-2P-Kv recordings are from a previous study^1^. n = 4 neurons for each GEVI. (**I**) F1 scores for the corresponding precision-recall curves shown in panel H, the horizontal black line denotes mean. P-value from an unpaired t-test. (**J-O**) 2P resonant-scan imaging of spontaneous activity in awake, behaving mice. (**J**) Experiment schematic. (**K**) Left: 2P image of a layer-5 FORCE1s-Kv-expressing neuron at the size and resolution used during voltage imaging. Right: Detrended trace of a 32-min, 400-Hz scan of the neuron shown. (**L**) Representative spiking events from panel K. (**M**) Spike rates. Acquisition frequency: 414-480Hz. Spike rates shown as a rolling mean computed using a 1-min window and 250-ms steps, and normalized to the mean firing rate in the first minute. n = 4 neurons from 3 mice. (**N**) Spike-triggered average from panel K. n = 1207 singlets. Horizontal line corresponds to the trace baseline average. (**O**) Comparison of spike response signal-to-noise ratios (SNR) between the start and end of recordings. n=4 neurons from 3 mice.

We next evaluated FORCE1s-Kv for extended 2P voltage imaging (without electrophysiology) in deep cortical layers in awake, behaving mice (Fig. 2J). Using continuous imaging of individual layer-5 neurons at 400 Hz for 30 minutes or more, optical spikes were detected across the recording period (Fig. 2K-L, M) with a response amplitude exceeding 100% dF/F (Fig. 2N; 4/4 neurons). Optical spike SNR remained high throughout the recordings (Fig. 2O).

Next, we compared FORCE1s-Kv to JEDI-2P-Kv in head-fixed, behaving mice using ultrafast local volume excitation (ULoVE) microscopy, leveraging this 2P technique’s capacity to assess GEVIs with supra-kilohertz temporal resolution and high sensitivity in single-cell recordings^4^. We chose JEDI-2P for comparisons, given that it is the indicator that produced the largest reported spike responses under ULoVE^5,9^. We conducted 10-min recordings of layer-2/3 cortical neurons at 7.1 kHz and depths of 117–303 µm (217 ± 55 µm, mean ± SD used here and henceforth; n = 26 cells from 4 mice).

Compared with JEDI-2P-Kv, FORCE1s-Kv generated responses to isolated spikes with a 3.4-fold increase in amplitude (92.7 ± 24.9 % vs. 27.6 ± 4.1%) (Fig. 3A-B). The slower off-kinetics of FORCE1s-Kv led to an accumulation of spike amplitudes, with bursts showing up to a 326% response (180 ± 90%, n=9 neurons; Fig. 3A, zoom-in epochs). FORCE1s-Kv also produced 1.8-fold larger responses than JEDI-2P-Kv to subthreshold fluctuations (40 ± 10 vs. 22 ± 4 %; Fig. 3C).

**Figure 3.**
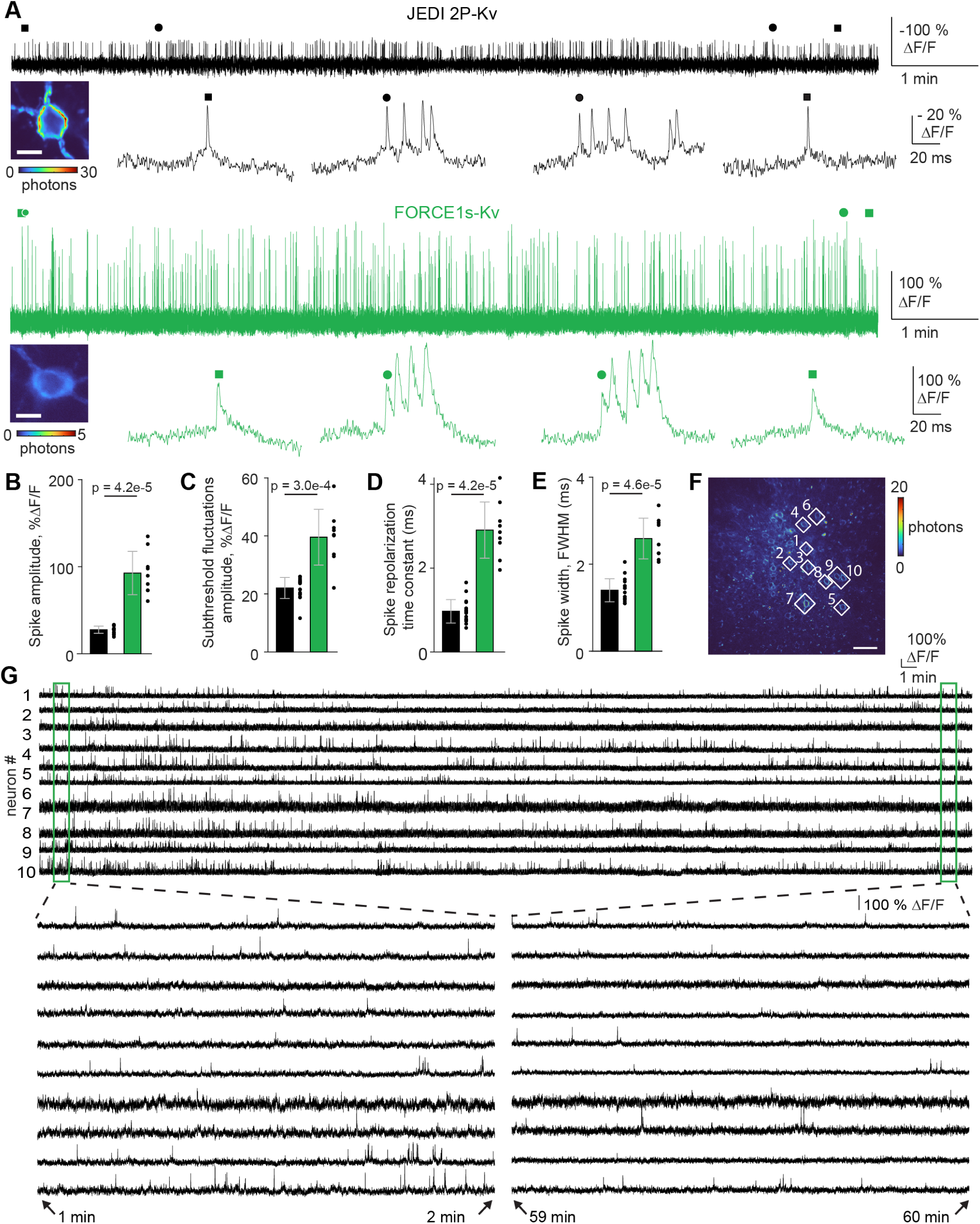
FORCE1s-Kv reports spikes and subthreshold fluctuations with larger amplitudes than JEDI-2P-Kv in vivo. All recordings were performed in layer 2/3 of visual cortex area V1. (**A**) Representative 10-min, 7.1-kHz ULoVE recordings of JEDI2P-Kv and FORCE1s-Kv. The recorded neurons are shown; scale bar: 10 µm. JEDI2P-Kv responses were inverted to facilitate comparisons with FORCE1s-Kv. Zoom-ins epochs show randomly picked isolated spikes (squares) and 4-spike bursts (circles) from the beginning and end of the recording. Note that the zoom-ins’ vertical scales differ by a factor of 5 between the two GEVIs. (**B-E**) Quantitative comparisons. Error bars: SD. n =17 (JEDI2P-Kv) and 9 (FORCE1s-Kv) cells. P-values from two-sample Wilcoxon rank sum tests. (**F**) Image showing 10 FORCE1s-Kv-expressing neurons selected for imaging. White lozenges: scanned areas. (**G**) 1-hr-long, 250-Hz simultaneous diagonal scan recordings of the 10 neurons from panel F. Zoomed-in epochs from the beginning and end of the recordings are shown.

FORCE1s-Kv exhibited repolarization kinetics of 2.9 ± 0.7 ms, ∼3-times slower than the 1.0 ± 0.3 ms of JEDI-2P-Kv (Fig. 3D). Consistently, FORCE1s-Kv reported spikes with an optical spike full width at half maximum of 2.6 ± 0.5 ms, 1.9-fold larger than those reported by JEDI-2P-Kv (1.4 ± 0.3 ms) (Fig. 3E).

Compared with JEDI2P-Kv, FORCE1s-Kv displayed a slower photobleaching rate during the initial phase (Fig. S3.1A, *left*), and a 1.7-fold lower slope in the power-law decay for the subsequent phase (Fig. S3.1A, *right*). Spike amplitude and width remained stable throughout the recordings for both GEVIs, suggesting that continuous illumination did not overtly impact the underlying electrical waveform or the optical properties of the indicators (Fig. S3.1B). Robust spiking activity was reported throughout the recording, with an excellent signal-to-noise ratio. Both indicators exhibited a modest reduction in firing rate, possibly reflecting decreased behavioral activity.

As expected from its inverted response polarity, the dim-to-bright FORCE1s-Kv’s exhibited a lower baseline photon flux than the bright-to-dim JEDI-2P-Kv’s (Fig. S3.1C). The fold-difference (3.3-fold) was slightly greater than that observed in vitro (1.9), reflecting differences in experimental conditions or confounding factors, such as differences in expression levels or virus preparation. Despite this lower observed baseline photon flux, spike SNR with FORCE1s-Kv still outperformed JEDI-2P-Kv by 1.8-fold (8.5 ± 1.9 vs 4.6 ± 1.2 Fig S3.1C). The larger response amplitude and slower off-kinetics of FORCE1s-Kv more than compensated for its lower baseline brightness, producing 3.0-fold greater detectability than JEDI-2P-Kv, as measured by the D’ metric^12^ (27.9 ± 8.1 vs 9.4 ± 2.7; Fig. S3.1C). Subthresholds’ SNR were comparable between the two sensors (FORCE1s-Kv: 3.7 ± 1.3; JEDI2P-Kv: 3.7 ± 0.9; Fig S3.1C).

### FORCE1s-Kv enables scalable, extended multi-cell voltage imaging

We asked whether the enhanced sensitivity and kinetics of FORCE1s tuned for sub-kilohertz sampling would enable multi-cell spike-resolved imaging. ULoVE optical recording requires sparse labeling to reduce SNR degradation caused by neuropil contamination. To overcome this constraint, we implemented a novel random-access “diagonal scanning” approach, using acousto-optic deflectors to rapidly acquire from multiple single-soma ROIs. Unlike ULoVE, diagonal scanning generates images, enabling the removal of background and neuropil pixels during analysis. It thereby achieves a higher effective SNR than holographic-shaped PSFs in densely labeled preparations where neuropil contamination is substantial^18^ (Morizet et al., manuscript in preparation).

We employed diagonal scanning at 210–250 Hz, as sampling below 200 Hz impaired detection of isolated spikes and individual action potentials within bursts (Figs. S3.2-3). At these sub-kilohertz sampling rates, FORCE1s-Kv supported one-hour recordings from 8–10 simultaneously imaged neurons in awake, behaving mice (n = 5 FOVs, 3 mice; Figs. 3F–G, S3.4–S3.8). These results show that FORCE1s-Kv enables multi-cell, sustained, and spike-resolved two-photon recordings using an acquisition regime that is compatible with widely used resonant-scanning 2P microscopes.

To image larger populations, we next used a 2P microscope equipped for FACED2.0, a modality previously shown to support voltage imaging with JEDI-2P-Kv^8^. However, JEDI-2P-Kv requires very high frame rates to maintain spike fidelity. When we downsampled JEDI-2P-Kv datasets from 800 to 400 and 200 Hz, spike z-scores declined moderately (Fig. S4.1A). However, spike counts dropped sharply (Fig. S4.1B), presumably because spike responses were more likely to occur partially or entirely outside the fluorescence sampling window.

For functional mapping, we used a 256 Hz frame rate—within the range that preserved FORCE1s-Kv’s spike detectability and aligned with the upper frequencies used for diagonal scanning. In awake, head-fixed mice viewing drifting gratings (0.5-s blank, then 0.5-s motion in one of eight directions), we imaged a 429 × 532 µm FOV in V1—∼3.6× larger than achieved with JEDI-2P-Kv at 800 Hz. We identified 154 somata (Fig. S4.2A), extracted single-trial traces, and computed trial-averaged subthreshold responses. Spike detection used a threshold targeting an estimated 1% false-positive rate (Fig. S4.2B). Spikes were clearly separable from subthreshold fluctuations in raw, single-trial traces (Figs. 4C–D), and subthreshold tuning was apparent within and across trials. FORCE1s-Kv demonstrated high photostability across 22 trials spanning 180 s, retaining 86 ± 6% of initial fluorescence (mean ± SD; Fig. 4B).

**Figure 4.**
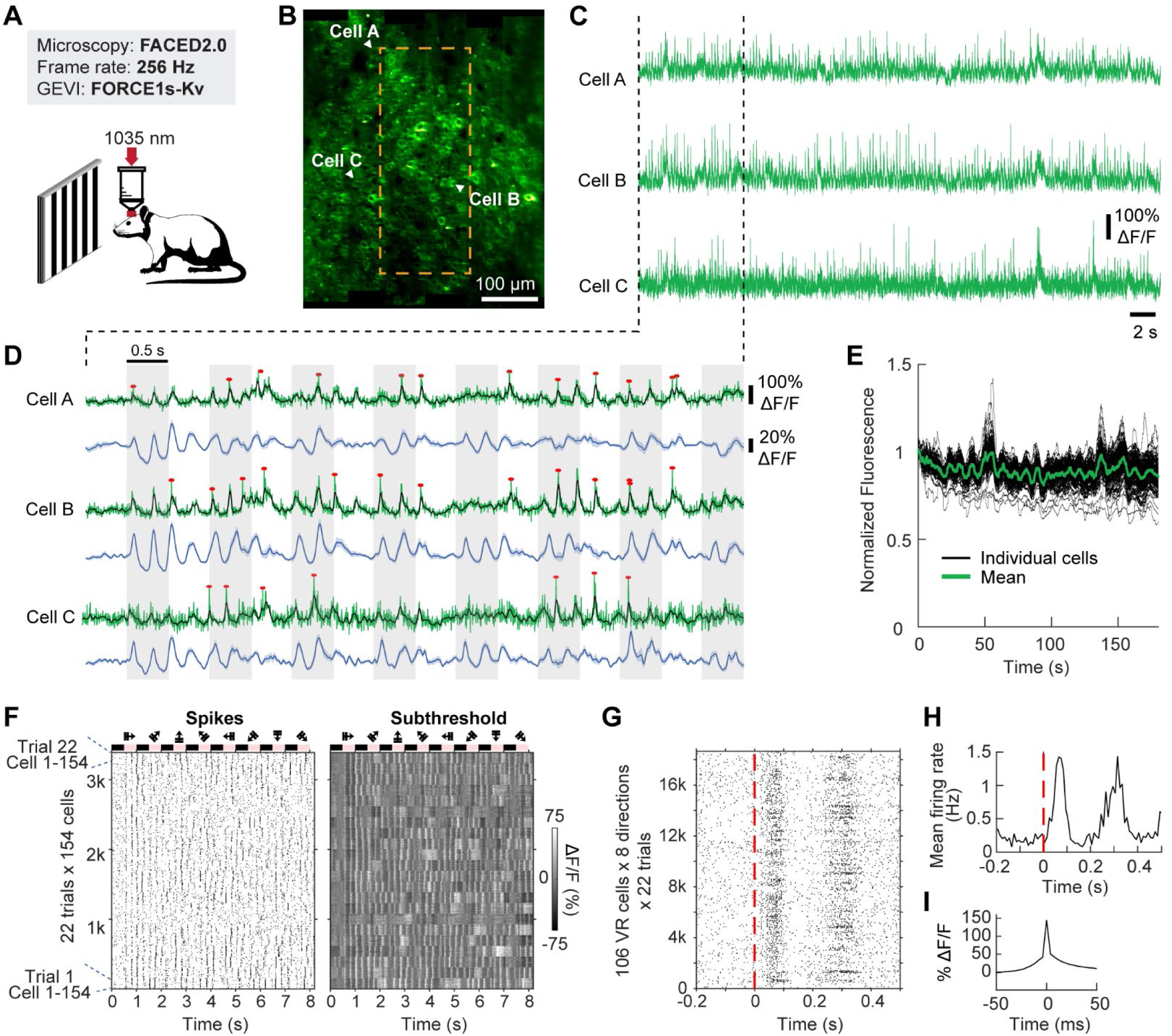
FORCE1s-Kv enables large-scale recording of voltage activity via FACED2.0 2P fluorescence microscopy. (**A**) Experimental schematic. Layer-2/3 neurons of visual cortex area V1 were imaged with FACED2.0 microscopy in head-fixed, awake mice during grating stimulation. (**B**) A representative 429×532-µm FOV of 154 neurons imaged at a frame rate of 256 Hz. The orange dashed box outlines the relative field-of-view (160×400 µm) used in earlier JEDI-2P recordings acquired at 800 Hz. (**C**) 41-s raw single-trial recordings from neurons labeled in panel B. (**D**) Zoomed-in epochs from panel C display raw responses (green) with low-pass-filtered subthreshold fluctuations (black) and detected spikes (red asterisks). Blue traces show trial-averaged subthreshold activity. Shaded areas: SEM. Gray-shaded blocks: periods with visual stimulation. (**E**) Visualization of GEVI photostability over 180 s. Black: traces from individual neurons, normalized to initial brightness and smoothed with a 4-second moving window. Green: mean. n = 154 neurons. (**F**) Raster plots of spikes (*left*) and subthreshold fluctuations (*right*) from all cells and trials, grouped by trial. Grating stimulus timing and orientation are shown above the plots. (**G**) Spike raster plot aligned to visual stimulus onset (red dashed line) from 106 visually responsive cells. (**H**) Mean firing rates of cells in panel F. Bin width: 10 ms. (**I**) Average spike profile. n = 10,853 spikes across all 154 cells.

Across the population, neurons displayed synchronized subthreshold dynamics with trial-to-trial variability (Fig. 4E). Out of the 154 neurons, 138 (89%) exhibited visually evoked subthreshold activity, and 106 (69%) were visually responsive (VR) based on spiking. Averaging over 18,656 1-s trials (106 VR neurons × 8 directions × 22 trials), spike rates increased by up to 8-fold during grating presentation (Figs. 4F–G). Averaging 10,853 detected spikes yielded a mean spike (including the underlying subthreshold) of 144% ΔF/F (Fig. 4I), compared with −20% for JEDI-2P-Kv, underscoring FORCE1s-Kv’s large positive-going responses.

We next tested whether FORCE1s could support multi-cell voltage imaging on a standard resonant-scanning microscope. Using this setup, FORCE1s enabled simultaneous imaging of six neurons at 400 Hz, yielding high-SNR spike and subthreshold responses (Fig. 5A–C). In contrast, JEDI-2P recordings under comparable conditions are typically restricted to single neurons¹, due to the indicator’s smaller response amplitude (Fig. 1J), lower detectability (Fig. 2I), fast repolarization kinetics, and negative-going design—which increases the need for sparse expression to avoid neuropil contamination. While we chose 400 Hz here due to FORCE1s’ high F1 score at this rate (Fig. 1J), our FACED and diagonal-scanning results indicate that 200–250 Hz imaging should also be feasible, enabling even larger neuron counts on resonant-scanning systems.

**Figure 5.**
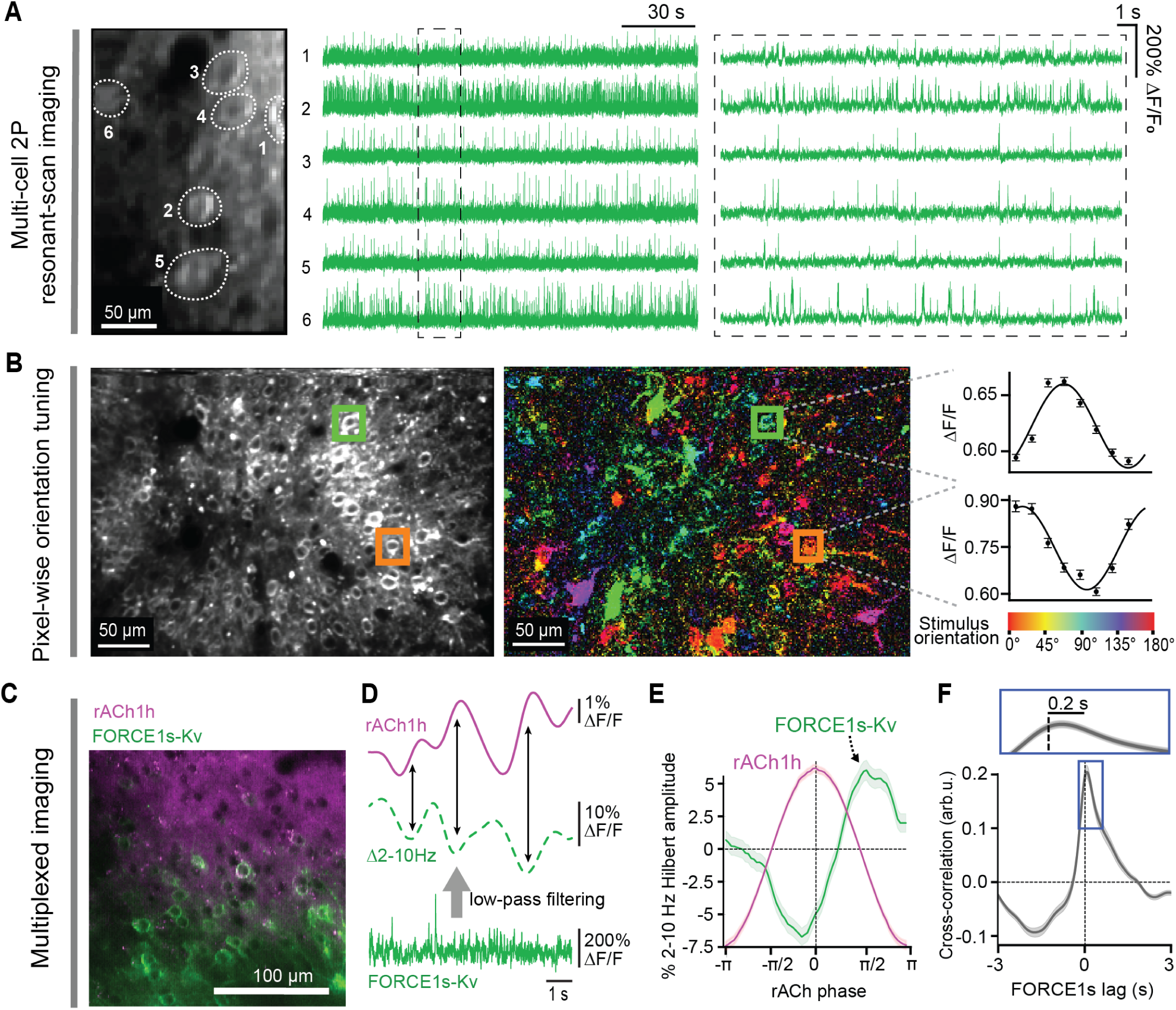
FORCE1s enables functional imaging and multi-modal integration. All data acquired with 2P resonant-scan imaging in visual cortex area V1, layer 2/3. (**A**) Simultaneous imaging of n = 6 neurons. Frame rate: 400 Hz. *Left*: outlines show the regions of interest. *Middle*: Unprocessed ΔF/F_0_ traces. *Right*: Zoomed-in epochs from the dashed box. (**B**) Pixel-wise orientation tuning. Frame rate: 115 Hz. *Left*: analyzed field-of-view. *Middle:* Pixelwise tuning map. Colors indicate the preferred orientation of the visual stimulus. *Right*, tuning curves for two outlined neurons. The color of the bounding box corresponds to their preferred orientation. Error bars: ± SEM (**C-F** Multiplexed voltage and acetylcholine imaging. (**C**) Field-of-view showing neurons expressing FORCE1s-Kv (green) and/or the acetylcholine sensor GRAB_rACh1h_, hereafter called rACh1h (violet). The two indicators were delivered via separate injections at nearby sites. (**D**) Simultaneous voltage and cholinergic imaging. Frame rate: 395 Hz. Arrows highlight how FORCE1s-Kv’s low-frequency activity is at a local minimum when acetylcholine levels are increasing. (**E**) GRAB_rACh1h_ phase-binned change in the low-frequency activity (2-10 Hz Hilbert amplitude) of FORCE1s-Kv fluorescence. Shaded areas: SEM. n=37 FORCE1s-kv expressing cells from one animal. (**F**) Mean cross-correlation between FORCE1s-Kv and GRAB_rACh1h_. Shaded areas: SEM. n=19 FORCE1s-Kv-expressing cells from one animal.

Finally, we evaluated FORCE1s’ ability to report neuronal functional properties across larger populations by quantifying stimulus selectivity in layer-2/3 visual cortical neurons—a standard benchmark in prior sensor evaluations^19,20^. The resonant-scanning frame rate was reduced to 115 Hz, enabling imaging of a 392 × 250 µm (200 × 314 pixels) field of view. We computed pixel-wise orientation tuning, which reveals cell-shaped patterns only when responses are robust and reliable, since single pixels are analyzed independently. FORCE1s-Kv-expressing neurons exhibited reliable pixel-level responses and diverse orientation preferences, consistent with the salt-and-pepper organization of the mouse visual cortex^21^ (Figs. 5B).

Overall, these results show that FORCE1s can increase the FOV size or the number of cells recorded using widely used (resonant scanning), cutting-edge (FACED2.0), and novel (diagonal scanning) methods.

### FORCE1s enables multi-modal interrogation of neural circuits

To address strong community interest in multiplexed recordings of neural activity, we evaluated whether FORCE1s can support simultaneous imaging of voltage and neurotransmitters. Both low-frequency membrane potential dynamics (Hilbert amplitude of 2–10 Hz oscillations) and ACh levels covary with pupil size, a well-established proxy for changes in arousal and attention^20^. However, concurrent measurements of voltage and cholinergic activity in awake, behaving mice were not readily attainable with previous methods, such as *in vivo* whole-cell patching, limiting efforts to examine their relative timing.

We co-expressed FORCE1s-Kv and a red-shifted acetylcholine sensor (GRAB_rACh1h_; hereafter, rACh1h) in the visual cortex (Fig. 5C). FORCE1s-Kv resolved changes in low-frequency activity across the rising and falling phases of cortical acetylcholine levels. These oscillations were dampened when ACh levels were high and became more prominent as ACh declined (Figs. 5D–E). Cross-correlation analysis revealed a peak slightly after zero lag, indicating that the FORCE1s-Kv signal tends to follow rACh1h maxima with a brief delay (Fig. 5F). Together, these results demonstrate that FORCE1s enables multimodal, brain-state-dependent activity measurements.

#### FORCE1s enables extended functional 2P voltage imaging in freely moving mice using MINI2P

2P imaging in freely moving animals promises deep-tissue, high-resolution readouts during naturalistic behavior, but applications have largely been confined to structural and calcium imaging^22–25^. Only a single study has reported voltage imaging in freely moving animals, and that work was restricted to brief (∼90-s) sessions, lacked functional validation, and relied on a non-commercial miniscope requiring a specialized 2-MHz laser—a configuration far beyond the reach of most laboratories^26^.

To determine whether FORCE1s could overcome these barriers, a miniature 2P microscope based on our MINI2P prototype^24^ —now commercially available— which we equipped with a slightly faster MEMS scanner (6.4 vs 5.6 kHz along the fast axis; Fig. 6A). FORCE1s-Kv was expressed in L2/3 V1 neurons via local viral delivery, producing moderately dense expression by 3–4 weeks (Fig. 6B). We first used the full 300 × 300 µm MINI2P FOV to locate neurons, then zoomed into 30–50 µm regions (1–3 neurons) for functional imaging (Fig. 6B). Reducing the imaging area (32–64 pixels × 24 lines) enabled frame rates up to 427 Hz, matching framerates that produced robust single-spike detection in resonance scanning (Fig. 2H, I). Voltage imaging was conducted while mice foraged freely in an 80 × 80-cm open field for ≥40 min per session, while behavior was tracked using an overhead IR camera (Fig. 6C).

**Figure 6.**
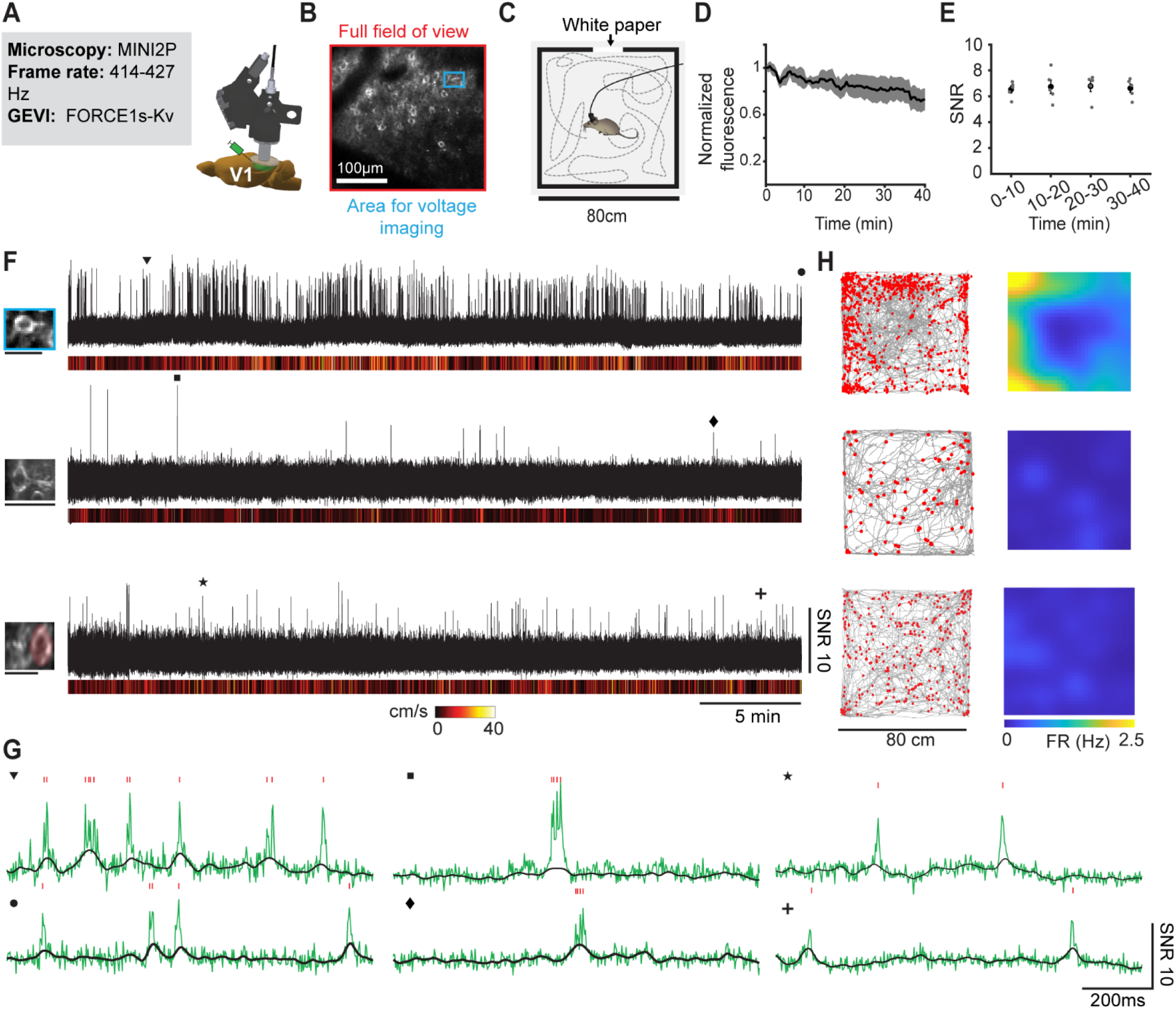
FORCE1s enables extended 2P voltage imaging in freely moving mice using MINI2P. (**A**) Imaging parameters and experimental schematics. FORCE1s-Kv was expressed in L2/3 neurons of cortical area V1. (**B**) Representative MINI2P-acquired average intensity projection image showing FORCE1s-Kv expressed in neuronal somata. To achieve a sampling rate > 400 Hz, functional imaging was restricted to fields of view of up to 70 × 30 μm, (e.g., blue box) covering 1–3 neurons. (**C**) Mice equipped were placed in an enclosed 80×80-cm open field arena and allowed to explore freely for up to 40 min. A white A4-size paper served as a visual cue. (**D**) FORCE1s-Kv photobleaching. Laser power: 50–65 mW, measured out of the objective. Mean baseline fluorescence is plotted. Shaded areas: SEM. n = 5 neurons from two mice. (**E**) Signal-to-noise ratio (SNR) is stable over 40-min recordings. SNR was quantified as the ratio of the optical spike amplitude to the fluorescence standard deviation. Same neurons as in panel D. SNR during the first 10 min: 6.2 ± 0.4; SNR during the last 10 min: 6.3 ± 0.5; P = 0.99 (Paired Wilcoxon Signed-Rank Test). n = 5 neurons from 2 mice. (**F**) 40-min recordings from three example neurons. The animal’s speed is shown below. Symbols indicate zoom-in epochs in panel G. Left: imaged neurons. The third trace corresponds to the neuron highlighted with the red overlay. Scale bars: 30 μm. (**G**) 1-s epochs from panel F. The overlaid red lines indicate subthreshold fluctuations. Vertical red ticks mark VolPy-detected spikes. (**H**) Spatial activity maps of the neurons shown in panel F. *Left*: animal trajectories (gray lines) with VolPy-detected spikes (red dots). *Right*: corresponding spatial firing rate maps. FR = firing rate.

Across 40-min sessions, MINI2P imaging of FORCE1s-Kv detected spike with high SNR of 6.29 ±0.49 (mean ± SEM, Fig. 6F). Single spikes were detected even during rapid locomotion (> 20 cm/s), indicating that fast voltage dynamics can be faithfully tracked under freely moving conditions. FORCE1s-Kv was photostable, retaining 72 ± 9% (mean ± SEM) of baseline fluorescence at session end (Fig. 6D), and SNR remained stable from start to finish (Fig. 6E).

To illustrate functional mapping capabilities, we constructed spatial firing-rate maps during exploration (Fig. 6G). One example neuron showed elevated firing near arena boundaries (border-tuning score 0.60 vs. shuffled controls 0.28 ± 0.18; Fig. 6F-G), indicating that FORCE1s can report meaningful, behaviorally relevant functional readouts in freely moving animals.

These findings demonstrate that sustained, single-spike–resolution functional two-photon voltage imaging is achievable in freely moving mice using FORCE1s combined with a cost-effective MINI2P system, comparable to commercially accessible offerings.

## Discussion

FORCE1s overcomes a central barrier to two-photon voltage imaging by coupling large positive-going spike responses (∼100% ΔF/F) with off-kinetics tuned for sub-kilohertz sampling. This combination enables spike-resolved, multi-cell recordings on standard resonant-scanning microscopes, the most common systems in calcium-imaging laboratories, thereby lowering technical and cost barriers to dissemination. FORCE1s further extends deep-tissue voltage imaging to freely moving two-photon miniscopes using accessible MEMS hardware, maintaining high SNR and strong photostability over prolonged sessions. Collectively, these capabilities will help transition 2P voltage imaging beyond a niche, platform-dependent technique to a robust, community-accessible approach for interrogating subthreshold dynamics and spiking activity during both head-fixed and naturalistic behaviors.

Beyond standard configurations, FORCE1s scales on ultrafast acquisition modalities. With diagonal scanning, it sustained hour-long recordings from groups of 8-10 user-selected neurons. With FACED2, it expanded the field of view by ∼3× while preserving spike fidelity—avoiding the spike-loss penalty that constrains faster, lower-response indicators.

FORCE1s also provides a tractable scaffold for future engineering. Priorities include faster on-kinetics and greater dynamic range to push recordings to deeper layers, smaller structures, and larger populations. FORCE1s’ deliberately slower off-kinetics, which preserve spike detection in 200-400-Hz multi-cell recordings, broaden the optical spike waveform. While advantageous to scaling up recordings, this widening is less well suited for measuring sub-millisecond spike timing, capturing AP rates in fast-spiking cells, and faithfully reconstructing voltage waveforms. Those use cases call for an ultrafast companion indicator, which we are developing in parallel, ensuring a complete kinetics toolkit for circuit studies.

In sum, FORCE1s advances both accessibility and scale in two-photon voltage imaging. By enabling routine, high-SNR, multi-cell recordings in behaving animals on standard hardware—and enlarging fields of view on ultrafast modalities—FORCE1s provides a versatile, robust tool for probing neuronal computation in behaving animals.

## Methods

### High-throughput GEVI screening

#### Predicted GEVI 3D structures and directed evolution lineage tree

Predicted 3D models of the FORCE1s sensors were generated by introducing targeted in silico mutations into a model of JEDI-2P generated by AlphaFold3, using the mutagenesis wizard of the PyMOL Molecular Graphics System (Version 3.0, Schrödinger, LLC). The evolutionary tree of mutations was rendered using Microsoft Office Visio.

#### Solutions

- **Growth medium #1**: is comprised of high-glucose Dulbecco’s Modified Eagle Medium (Sigma-Aldrich, D1145-500ML) supplemented with 10% fetal bovine serum (FBS, Sigma-Aldrich, F2442-500ML), 2 mM glutamine, 100 unit/mL penicillin, 100 μg/mL streptomycin, and 750 μg/mL of the antibiotic G418 sulfate (geneticin).
- **Growth medium #2**: is comprised of high-glucose Dulbecco’s Modified Eagle Medium supplemented with 5% FBS, 2 mM glutamine, 100 U/mL Penicillin, and 100 μg/mL Streptomycin.
- **Imaging solution #1**: is comprised of 110 mM NaCl, 26 mM sucrose, 23 mM glucose, 20 mM HEPES, 5 mM KCl, 2.5 mM CaCl_2_, 1.3 mM MgSO_4_, adjusted to pH 7.4 with NaOH, and adjusted to 300 mOsm/kg with H_2_O
- **Imaging solution #2**: is comprised of 106.5 mM NaCl, 26 mM sucrose, 23 mM glucose, 20 mM HEPES, 8.5 mM KCl, 2.5 mM CaCl_2_, 1.3 mM MgSO_4_, adjusted to pH 7.4 with NaOH, and adjusted to 300 mOsm/kg with H_2_O
- **Internal solution:** is comprised of 115 mM K-gluconate, 10 mM HEPES, 10 mM EGTA, 10 mM glucose, 8 mM KCl, 5 mM MgCl_2_, 1 mM CaCl_2_, adjusted to pH 7.4 with KOH, and adjusted to 290 mOsm/kg with H_2_O

#### Library construction

GEVI libraries were generated by site-directed saturation mutagenesis targeting single residues. To obtain a more uniform distribution of residues for saturation mutagenesis, we combined primers containing the NNT, VAA, ATG, or TGG codons (N = any base; V = A, G, or C) at a molar ratio of 16:3:1:1. The 20-μL PCR reaction mix contained 1 μL of a 10-μM forward primer mix, 1 μL of a 10-μM reverse primer, 5 ng of template plasmid, and 10 μL of a 2× PCR master premix (PrimeSTAR HS DNA polymerase, Takara). DNA was amplified using the following protocol: an initial denaturation step at 98°C for 30 s; 30 amplification cycles consisting of 98°C for 10 s, 57°C for 10 s, and 72°C for 1 minute/kb of fragment length; and a final extension step at 72°C for 5 minutes. The pcDNA3.1/Puro-CAG^27^ backbone was linearized using the restriction enzymes NheI and HindIII. PCR products and linearized backbones were purified using gel electrophoresis and the GeneJET Gel Extraction Kit (Thermo Fisher Scientific). PCR products, including a C-terminus GSSGSSGSS-linked^28^ mBeRFP^29^ E6F reference protein, were assembled in the vector backbone using the In-Fusion assembly system (In-fusion HD Cloning Plus, Takara) according to the manufacturer’s instructions. The mBeRFP CDS used for amplification had an E6F mutation compared with the original sequence; this variant of mBeRFP was used for all cloning of mBeRFP. The In-Fusion reaction mix was transformed into commercial chemically competent bacteria (XL10-Gold, Agilent) with a transformation efficiency exceeding 5×109 FU per μg of DNA. Liquid cultures were inoculated with manually picked colonies, and purified plasmids were prepared using a 96-well plasmid purification kit (96-well Mini Plus Plasmid Extraction System, Viogene #GF961001) following the manufacturer’s instructions.

#### In vitro characterization plasmid construction

Plasmids used for in vitro characterization were assembled using the In-Fusion cloning technique (Takara Bio USA, Inc.) into a pcDNA3.1/Puro-CAG vector with or without a C-terminus GSSGSSGSS-linked mBeRFP reference protein^5^. Liquid cultures were inoculated with manually picked colonies, and purified plasmids were prepared using a purification kit (Mini Plus Plasmid Extraction System, Viogene #GF2002) following the manufacturer’s instructions. The plasmid sequence was confirmed using whole-plasmid nanopore sequencing (Plasmidsaurus Inc.) or Sanger sequencing (Eurofins Genomics LLC). The ASAP4e sequence was amplified from the pAAV-hsyn-ASAP4e-Kv-WPRE plasmid (Addgene, plasmid # 201031). The pcDNA3.1/Puro-CAG-EGFP plasmid was generated in our lab previously^5^.

#### Cell culture and transfection in G6-well plates

To screen GEVIs for responses within the physiological range, we used HEK293 cells stably expressing the human Kir2.1 channel^30^, which maintains a resting membrane potential of approximately −83 mV using imaging solution #1. This membrane potential was chosen because it approximates the lower bound of the physiological range for mammalian neurons, enabling screening for improved responses across the hyperpolarized state and into the depolarized range.

HEK293-Kir2.1 cells were cultured at 37°C with 5% CO2 in growth medium #1. G418 was added to maintain the expression of the Kir2.1 transgene, which was chromosomally integrated with a G418 resistance gene. For screening GEVIs, glass-bottom 96-well plates (P96-1.5H-N, Cellvis) were first coated with poly-D-lysine (30-70 kDa) to promote cell adherence to the glass. The coating was performed for 1 hour at room temperature or 37°C, and the plates were washed twice with Dulbecco’s Phosphate Buffered Saline (Corning, #21-031-CV). HEK293-Kir2.1 cells were then plated to 60-80% confluency in 100 μL of growth medium #2.

We randomly selected 88 variants per saturation mutagenesis library. According to statistical modeling^31^, our library generation and sampling strategy produced a ∼99% theoretical probability that any given screened library included the best residue. Each well was transfected with jetPrime (Polyplus) according to the manufacturer’s protocol, using the following specifications: a mixture of 130 ng DNA, 0.4 μL jetPRIME transfection reagent, and 10 μL jetPRIME buffer was mixed with 50 μL of growth medium #2, and then added to the well. Independent transfections were defined as those in which DNA was added to each well separately. After four hours or the next day, 100 μL growth medium #2 from each well was replaced with fresh growth medium #2 to minimize potential cytotoxicity from the transfection reagents. Two days post-transfection, the cells were washed twice with 150 μL of imaging solution #1 at room temperature. Wells were filled with 100 μL of the imaging solution #1 and screened with our two-photon high-throughput screening platform (see below). The imaging solution #1 was adjusted with H2O to achieve a final osmolarity of 290-310 mOsm/kg.

#### Two-photon GEVI screening system

Two-photon library screening was completed using roughly the same hardware as previously described^5^. An inverted microscope with multi-photon capability (A1R-MP, Nikon Instruments) was used for two-photon screening and in vitro characterization of GEVIs. The two-photon excitation light was generated by a Ti:Sapphire femtosecond laser (Chameleon Ultra II, Coherent) with a repetition rate of 80 MHz and a tuning range of 680 nm to 1,080 nm. Laser power was set using an acousto-optic modulator and delivered to the sample plane through a 20×0.75-NA objective (CFI Plan Apochromat Lambda, Nikon Instruments). The emission light from the sample was split using a 560-nm dichroic mirror (348958, Chroma) and filtered by 525/50-nm (353716, Chroma) and 605/70-nm filters (Nikon Instruments) for green and red channels, respectively. Emitted photons were detected by gallium arsenide phosphide (GaAsP) photomultiplier tubes (PMTs). A motorized travel stage (H139E1, Prior or SCANplus IM 130×85 −2mm, Märzhäuser Wetzlar GmbH & Co. KG) was used to control the position of the field of view, hold 96-well plates during screening, and hold the perfusion chamber during electrophysiological characterization.

To support automation of the system, data acquisition and output boards (PCI-6229 and PCI-6723, National Instruments) were connected to the microscope computer through a PXI Chassis (PXI-1033, National Instruments). The computer was equipped with an Intel Xeon Gold 6334 Processor, 8-core, 3.60 GHz, 256 GB of DDR4 RAM, and two 12TB 7200 RPM hard drives configured in RAID 0 to facilitate high-speed imaging. JOBs scripts in NIS-Elements HC (version 5.42.07, Nikon Instruments) were used to control the microscope system (e.g., stage position), manage the optical configurations (e.g., excitations), initiate image acquisition, and trigger the stimulator.

A digital isolated high-power stimulator (4100, A-M System) was used to provide electric field stimulation. Electric pulses were passed through a pair of electrodes made from 0.5 mm-wide platinum wires (99.95% pure, AA10286BU, Fisher Scientific). The two L-shaped electrodes, each 2 mm in horizontal length and 3 mm apart, were secured on a 3D-printed polylactic acid holder. The holder was fixed to a motorized linear translation stage (LSM050B-E03T4A-MC10T3, Zaber), which was used to raise and lower the electrodes in and out of individual wells. Two smaller manual linear translation stages (411-05S, Newport) were used to fine-tune the x-y position of the electrodes in the plane parallel to the screening plate. During electric field stimulation, the electrodes were submerged under the imaging solution #1, with the centers of the electrode wires 500-600 μm above the bottom of the well.

#### Two-photon GEVI screening

For each well of the screening plate, four non-overlapping fields of view (FOV) of 512 × 32 pixels were imaged per well at 440 Hz using a resonant galvanometer scanner. The laser was tuned to 940 nm and set to 34-50 mW at the sample plane. To collect images for the brightness calculation, fifty frames of the GEVI and the reference protein mBeRFP were collected in parallel on the green and red channels, respectively. To the same FOV, electric field stimulation was performed during continuous imaging of the green channel for 4000 frames or ∼9 s. All stimulation trains were generated in AMS4000 software (A-M Systems) and uploaded to the stimulator as a series of monophasic square pulses. The stimulation protocol started with five monophasic square pulses (60-V 1-ms) spaced 300-ms apart. This was followed by series of seven pulses spaced 300-ms apart: 15-V 1-ms, 25-V 1-ms, 30-V 1.5-ms, 30-V 2.5-ms, 30-V 3.5-ms, 30-V 4.5-ms, 30-V 7.5-ms. Next in the train was a 100-Hz train (10 monophasic square pulses with 2.5-ms width, 30-V amplitude, and an inter-pulse duration of 10 ms), and finally another five 60-V 1-ms pulses.

#### Analysis of high-throughput screening data

Image analyses were performed by custom routines in MATLAB (version R2024a, MathWorks) as previously described^5^. Time-lapse images recorded in ND2 format were imported into MATLAB using the Bio-Formats toolbox (version 6.3.1)^32^. For both channels (red and green) of each FOV, saturated pixels from any point during the videos (e.g., from over-expressing cells) were removed. Background correction was computed by subtracting the mode intensity of the first frame’s pixel intensity histogram from each non-saturated pixel’s value. An initial foreground mask was computed from the first 20 frames of each channel by applying pre-defined intensity thresholds for the green and red channels to distinguish pixels containing GEVI and reference protein signal from pixels just containing autofluorescence noise. Relative GEVI brightness (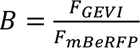, arb. unit) was calculated using the mean fluorescence intensity of the first 20 frames in the green channel (*F*_*GEVI*_), divided by the average fluorescence intensity of the first 20 frames in the red channel (*F*_*mBeREF*_). Relative GEVI brightness is reported to correct for FOV-to-FOV differences in the number of transfected cells, the number of selected pixels, and overall expression (e.g., due to pipetting errors or biological variation).

Quantification of response amplitudes from the 9 s recording could be distorted by the inclusion of non-plasma-membrane-localized GEVIs within an FOV that were bright but non-responsive. To account for non-responsive pixels, we applied an additional correlation-based selection step. Specifically, a template was created from the mean temporal trace of all remaining pixels in the recording. The temporal traces of each remaining pixel were correlated with the template using Pearson correlation coefficients. Pixels were ranked according to their correlation coefficients. To determine the optimal pixels, we evaluated the signal-to-noise ratio (SNR) across cumulative groups of pixels and identified the group corresponding to the peak SNR value. This subset was then used as the final output of the correlation filtering process. FOVs with final masks containing fewer than 300 pixels were removed from further analysis as reduced pixel selection often is representative of a low cell confluency, poor transfection, or a detrimental mutation. Using the pixels from the final mask, we calculated the total signal at each time point to measure fluorescence over time. We corrected the green (GEVI) fluorescence trace for photobleaching by dividing each trace by the best-fit three-term exponential on the mean fluorescence using data outside of stimulations. To quantify photostability, (*P*, arb. unit), the area under the curve (AUC) was calculated from the fluorescence vs time trace normalized to 1 at *t* = 0. The peak response amplitude (R) was quantified from the photobleaching-corrected fluorescence trace 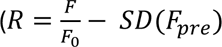, arb. unit), where *F* is the peak fluorescence, *F* is the mean baseline fluorescence for the first 100 ms of the recording, and *F*_*pre*_ is the baseline fluorescence for the 100 ms preceding the pulse. The standard deviation (SD) of *F*_*pre*_ was subtracted from 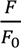 (response) to account for baseline fluctuations. Given that FOVs can vary in the number of pixels selected, the FOV area-weighted mean was taken for each well when determining the average response.

Compound metrics that consider multiple performance metrics were used to simplify the ranking of indicators^5^. We previously developed the detectability index metric 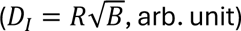 — where *R* is the peak response amplitude and *B* is the relative brightness. We also sought to consider the impact of photobleaching on voltage recording and avoid variants that are bright but bleach rapidly. Thus, we used the detectability budget 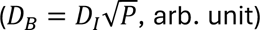 — which combines all measured performance characteristics by replacing the initial relative brightness in the *D*_*I*_ equation with the average relative brightness measured during the screening experiment. None of the above metrics include physical units, as they are relative parameters intended solely for comparative ranking of indicators during the screening process.

### GEVI characterization in vitro

#### Two-photon excitation spectra

HEK293A cells were cultured and transfected in 96-well plates with either FORCE1s, ASAP4e, SpikeyGi2, or JEDI-2P as discussed for HEK293-Kir2.1 cells in the “Cell culture and transfection in 96-well plates” section, excluding the use of G418 sulfate. HEK293A cells were used as they have a more depolarized resting membrane potential than HEK293 Kir2.1 cells, causing dim-to-bright sensors like FORCE1s to be brighter for characterization. Cells were washed and imaged in imaging solution #1, resulting in a resting membrane potential of −20.0 ± 0.3 mV (mean ± SEM, n = 4 HEK293 kir2.1 cells). Images were acquired using the same microscope as described in the “Two-photon screening system” section. Laser pulses were not pre-compensated for dispersion in the microscope optical path. Excitation wavelengths from 700 to 1080 nm were used in 10-nm increments and set to a power of 29-32 mW at the sample plane, as measured by a microscope slide power sensor (S170C Thorlabs). Each 512×512-pixel FOV was imaged at each wavelength using two galvanometer optical scanners set to acquire 1.1 s images. Each well had two FOV one for even wavelengths and one for odd wavelengths.

Analysis was done separately for even and odd images. Images were background corrected by subtracting the mode intensity of the first frame’s pixel intensity histogram, excluding saturated pixels. A maximum intensity projection, excluding saturating pixels, across all images was used to determine values for foreground segmentation using a user-defined threshold to optimize cellular masking for all constructs. The mean intensity of the foreground pixels was calculated for each wavelength. Slight deviations in the power between wavelengths were corrected by assuming a quadratic dependence of fluorescence on illumination power at the sample plane. All mean intensity values for a single FOV were then normalized to the mean intensity value of the first 920nm image. The even and the odd FOV data for each well were combined, and the whole series of images for a well was normalized to the maximum fluorescence value in the series.

#### One-photon excitation and emission spectra characterization

To characterize the one-photon excitation and emission spectra of GEVIs, we used the same vector used for in vitro GEVI characterization with whole-cell voltage clamp, i.e., pcDNA3.1/Puro-CAG-*GEVI*, and a control vector pcDNA3.1/Puro-CAG-EGFP. Transfections were performed in 10-cm tissue culture dishes (353003, Corning), and HEK293A cells were seeded one day before forward transfection at 60–65% confluency, equivalent to 720,000–780,000 cells per well or 5,280,000–5,720,000 cells per dish in growth medium #2. Transfections were performed 36–48 h before imaging using JetPRIME. The transfection medium was replaced 4 h after transfection with fresh growth medium #2 to minimize potential cytotoxicity from the transfection reagent.

On the day of the experiment, cells from a 10-cm dish transfected with the same construct were detached with trypsin, washed twice, diluted into imaging solution #1, and evenly split into three wells of a glass-bottomed 96-well plate (P96-1.5H-N, Cellvis). Pooling the cells into a dense preparation was essential to produce a strong signal that the plate reader could robustly detect. Untransfected cells were prepared with the same method and deposited in a separate well to determine the background autofluorescence levels. A handheld automated cell counter (Scepter 3.0, Millipore) was used to plate a similar number of cells between conditions.

Spectra were determined using a plate reader (Cytation 5, BioTek) to quantify fluorescence from wells of the 96-well plates prepared above. Emission spectra were acquired by exciting at 430/10 nm and measuring emitted photons from 460 to 650 nm in 1-nm increments, with a bandwidth of 10 nm. Individual spectral scans were corrected for autofluorescence by subtracting the values from untransfected cells at each wavelength and then normalized to their respective peaks. The final emission spectra were averaged from the normalized spectra of each scan. The peak emission wavelengths were averaged from the peak excitation/emission wavelengths of each scan.

#### Brightness under two-photon illumination

To determine the brightness under 2P illumination, HEK293-Kir2.1 cells were transfected with pcDNA3.1/Puro-CAG-*GEVI*-(GSS)_3_-mBeRFP plasmids and prepared for imaging, as described in “Cell culture and transfection in 96-well plates.” However, imaging solution #2 was used instead of imaging solution #1 to obtain a resting membrane potential of ∼-70 mV, approximating the resting voltage of mammalian cortical pyramidal neurons. Images of the cells were acquired using the same microscope as used for screening (see “Two-photon screening system”) with a 20× NA 0.75 objective (CFI Plan Apochromat Lambda, Nikon). Cells were excited with 34.13 mW of 940 nm light. Four 1024 x 1024 pixel FOVs were captured for each well using two galvanometer optical scanners with a dwell time of 2.4 µs. Green (GEVI) and orange (mBeRFP) emitted light was collected simultaneously by two PMTs, spectrally separated by a 560 LP dichroic mirror and either a 525/50 (GEVI) or 605/70 (mBeRFP) emission filter. Images were background-subtracted, and then the relative brightness was calculated from the foreground mask as the average pixel intensity of the GEVI channel divided by the average pixel intensity of the mBeRFP channel.

#### Photostability under two-photon illumination

Comparison of indicators’ photostability under two-photon excitation was carried out using the same protocol as described previously1 and briefly described below:

We used the same titanium:sapphire femtosecond laser (Chameleon Ultra II, Coherent) as described in Evaluating indicators’ response amplitude under two-photon illumination. HEK-Kir2.1 cells were seeded and transfected with pcDNA3.1/Puro-CAG-GEVI vectors as described in High-throughput GEVI screening under widefield one-photon illumination.

Two days after transfection, the medium in the 96-well plate was removed, and the cells in each well were washed with 200 µL of imaging solution 2 (see “Solutions”) twice. Then, 100 µL of imaging solution 2 was added to each well. Excitation laser was generated from a titanium:sapphire femtosecond laser (Chameleon Ultra II, Coherent) and directed to the sample plane by the resonant galvanometer scanners through the same 20× 0.75 NA objective (CFI Plan Apochromat Lambda, Nikon Instruments). Photobleaching was conducted under 940-nm (other GEVIs) excitation with a power of 40, 80, or 100 mW at the sample plane. For each well, four non-overlapping FOVs of 512 × 32 pixels were continuously imaged at 440 Hz for 6 minutes. The emission light from the cell was split using a 560-nm dichroic mirror (348958, Chroma), filtered by a 525/50-nm bandpass filter (353716, Chroma) and collected by a gallium arsenide phosphide (GaAsP) photomultiplier tube (PMT). Fluorescence traces were normalized to the first frame.

To quantify GEVI photostability under 2P, every 10th image from the video was binned. Foreground segmentation was performed, followed by background correction using a user-defined threshold to optimize cellular masking across all constructs, as described in the “Analysis of high-throughput screening data” section. Saturating pixels were also removed from further analysis. The mean fluorescence from the foreground pixels was normalized to the fluorescence of the first frame of the binned video. The AUC of the normalized photobleaching traces, over the entire (6.05 min) time courses, was computed using the trapezoid rule (GraphPad Prism). The AUC provides a robust, model-independent measure of photostability across illumination power levels. This approach avoids the complications of fitting photobleaching traces, which often require multiple exponential components whose number and relative weights can vary with illumination intensity, thereby complicating statistical analysis. To facilitate comparison, AUC values were normalized to the maximal AUC possible for that recording (i.e. AUC for a straight line of y=1 for the same duration of the recording), such that each data point represents a GEVI’s integrated fluorescence intensity as a percentage of the total signal that would be produced by an ideal, non-bleaching (infinitely photostable) GEVI. AUCs were calculated from n = 3-4 independent transfections per power level. The Kruskal-Wallis test, followed by a Dunn’s multiple comparisons test, was used to determine if the relationship between “integrated fluorescent intensity vs. power” differed among GEVIs. The fitting of the 37 mW fluorescence traces was performed using a double exponential equation that modeled a two-phase decay.

#### Whole-cell voltage clamp

HEK293A cells (Thermo Fisher Scientific) were plated on 30-70 kDa poly-D-lysine-coated circular cover glass (12 mm, #0, 633009, Carolina) at 30% confluence in growth medium #2, two days prior to imaging. Chemical transfection was performed on the same day as plating, using 100 ng of DNA and 0.3 μL of FuGENE HD (Promega Corporation) transfection reagent per well of a 24-well plate (Costar, #3524), following the manufacturer’s instructions. Unless indicated otherwise, pcDNA3.1/Puro-CAG plasmids expressing the GEVI with no reference protein were used. The cells were cultured at 37°C with 5% CO_2_ before and after transfection. The next day, the transfection media was replaced with fresh growth medium #2 to minimize potential cytotoxicity from the transfection reagent.

Glass micropipettes (1B150-F-4, World Precision Instruments) were prepared using a pipette puller (P1000, Sutter), achieving a tip resistance of 2-6 MΩ. Micropipettes were loaded with internal solution. The internal solution was adjusted with H_2_O to be 10 mOsm/kg lower than the imaging solution #1, which typically resulted in an internal solution between 280 and 300 mOsm/kg. The micropipette was mounted on a patch-clamp headstage (CV-7B, Molecular Devices) and positioned using a micromanipulator (SMX series, Sensapex or a GmbH Mini23, Luigs & Neumann). The coverslip seeded with the transfected cells was placed in a custom glass-bottom chamber, designed based on the Chamlide EC (Live Cell Instrument), with the bottom of the chamber made from a 24×24 mm #1 glass coverslip (Thermo Scientific). Cells were continuously perfused with an imaging solution #1 at approximately 4 mL/minute using a peristaltic pump (505DU, Watson Marlow). Whole-cell voltage clamp was achieved using a MultiClamp 700B amplifier (Molecular Devices). Patch-clamp data were recorded using an Axon Digidata 1550B1 Low Noise system with HumSilencer (Molecular Devices). Command voltage waveforms were compensated for the liquid junction potential of −11 mV. Recordings were considered satisfactory and included in the final analysis only if, both before and after the recording, the patched cell had an access resistance (*Ra*) < 20 MΩ and a membrane resistance (*Rm*) > 10 × *Ra*.

#### Whole-cell current clamp

The setup for the current clamp was identical to that of the voltage clamp (described above), with slight modifications. The resting membrane potential of HEK293-Kir2.1 cells was determined using a whole-cell current clamp mode in imaging solution #1 or imaging solution #2 and compensated by −11 mV to account for the liquid junction potential. HEK293-Kir2.1 cells were plated the day before patching.

#### Resting membrane potential characterization

Cells were plated on poly-D-lysine-coated glass coverslips two days before recording without being transfected. Cells were washed twice with imaging solution, and then fresh imaging solution was added to the recording chamber immediately before recording. During experiments, the bath solution was not perfused.

Cells were visualized under bright-field illumination and whole-cell patch-clamp recordings were obtained using a patch amplifier in zero-current (I = 0) mode. After establishing the whole-cell configuration (break-in), the resting membrane potential (RMP) was measured directly from the amplifier software (Multiclamp 700B) (I = 0) without additional series resistance or capacitance compensation. All recorded RMP values were corrected for the liquid-junction potential (−11 mV) computed for the bath and pipette solutions. Reported RMPs represent junction-potential–corrected values.

#### Simultaneous electrophysiology and two-photon illumination recordings

The same imaging hardware setup was used as described in the “Two-photon screening system” section, without using any filter cubes before the PMT. An oil-immersion 40× NA 1.3 objective (CFI Plan Fluor DIC H/N2, Nikon Instruments) was used to focus 940 nm light set to ∼16-44 mW onto the sample plane. Videos were taken with a resolution of 512×16 pixels and a frame rate of 721 Hz under resonant scanning microscopy. Recordings where noise was indistinguishable from the signal due to the low brightness of the cell were removed from the final analysis. Electrophysiological recordings of AP waveforms were done at ∼33°C, and cells were held at –70 mV. We clamped HEK293A cells to a variety of typical AP waveforms and monitored the resulting changes in GEVI fluorescence. We used an AP waveform that was recorded from a representative hippocampal neuron and modified it to have an amplitude of 100 mV and a full width at half maximum (FWHM) of 2 ms, to mimic the shape of layer 2/3 cortical neurons at room temperature.^33^ To further improve the probability of detecting a response to the AP waveform for sensors with slower on kinetics, we also included APs widened to 4 ms FWHM. Since all sensors easily detected the 2-ms FWHM we did not further characterize the 4-ms FWHM AP waveforms. Cells were stimulated with 5 AP waveforms with a FWHM of 2 ms at 2 Hz, 5 AP waveforms with a FWHM of 4 ms at 2 Hz and 10 AP waveforms with a FWHM of 2 ms at 100 Hz. In the same protocol, we also recorded GEVI response to a modified burst of APs recorded from adult mouse somatosensory cortex layer 5 pyramidal neurons, designed to mimic APs on top of subthreshold depolarizations. The spike burst exhibited a subthreshold depolarization of ∼24 mV (from a baseline voltage of −70 mV) and APs with amplitudes of 60–90 mV (from a subthreshold voltage of −56 mV). The APs in the spike burst are 1.5–2 ms FWHM.

To characterize the sensors’ fluorescence vs voltage curves under 2PM, cells were prepared as described above for voltage clamping at 33°C. The cells were simultaneously patched and illuminated with ∼20 mW of 940 nm light at the sample plane through the same 40x oil objective. Cells were held at −70 mV for 4 seconds, followed by a series of 1-second voltage steps (90, 70, 50, 30, 20, 0, −20, −40, −60, −80, −100, and −120 mV), each separated by 1 second at −70 mV. To reduce the amount of light to which the cells were exposed, a 64 x 64-pixel galvo-galvo scanning image (5.2 µs dwell time, 1.24 µm pixel size) was recorded every second for the duration of the patch clamp protocol.

#### Optical responses to spike waveforms

To quantify spike characteristics in vitro, cells were analyzed using the same in-house Python software. When a whole cell was present in the FOV, segmentation was performed using Cellpose 3. For partial cells, Otsu thresholding was applied within a manually drawn region of interest (ROI) to distinguish foreground from background pixels. A background ROI was also manually defined for each cell to allow for background subtraction. Pixels were included in the analysis only if their fluorescence intensity exceeded the background signal by at least two times for at least 80% of the frames, and no pixel reached saturation at any point during acquisition. Photobleaching correction was performed by isolating non-stimulated segments of the recording and fitting single-, double-, and triple-exponential decay models to the remaining fluorescence signal. The best-fit model was selected based on the highest R², the lowest root mean squared error, and the mean absolute error, while also ensuring that the model parameters corresponded to a local minimum in both the Bayesian Information Criterion (BIC) and the Akaike Information Criterion (AIC) to avoid overfitting. The selected model was then used to correct for photobleaching by dividing the intensity of each frame by the normalized bleaching curve. ΔF/F₀ was subsequently calculated from the background- and bleach-corrected fluorescence.

#### Voltage response curves

Cells were analyzed using in-house-developed custom Python software. Individual cells were segmented in each field of view (FOV) using Cellpose 3^34^. To isolate the cell edge, a Laplacian filter was applied to the binarized Cellpose3 mask. Pixels were excluded if they became saturated in any frame during the experiment or if their baseline fluorescence was less than 1.5 times the median background signal. Background fluorescence was defined as the signal from regions of the image that did not contain cells. Due to the relatively small number of frames and their wide temporal spacing, photobleaching was approximated using a linear regression fit to the fluorescence decay of frames acquired at a membrane potential of −70 mV. Photobleaching was corrected by multiplying the fluorescence time series by the inverse of the fitted bleaching decay function. ΔF/F₀ was calculated using the background and bleach-corrected signal, with F₀ defined as the average fluorescence intensity at −70 mV of the corrected signal. We then fit the calculated ΔF/F₀ to the logistic sigmoid 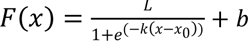 to determine the ΔF/F₀ at −70 mV which we then sued to translate out calculated ΔF/F₀ such that the fit function would have F(−70 mV) = 0 ΔF/F₀.

#### Kinetics

Kinetics were evaluated under one-photon illumination. The same imaging hardware setup described in the “Two-photon screening system” section was used, with the following modifications. Cells were illuminated with 475-nm light (SpectraIII L Light Engine, Lumencor) and conditioned using the 474/28-nm band of a multi-band dichroic mirror (77015970, Semrock). The irradiance at the sample plane was ∼4 mW/mm^2^. Green-emitted photons were reflected towards a multialkali photomultiplier tube (PMT, PMM02, Thorlabs) installed on one of the side ports of the microscope using the 493-nm band of the multi-pass dichroic (77015970, Semrock) and then filtered at 515/30-nm using a multi-pass filter (77015970, Semrock). Kinetics were evaluated using three 1-s 100-mV depolarization pulses from −70 to 30 mV. Between each pulse, cells were held at –70 mV for 1.4 s. Recordings were performed at 32-33°C using a feedback-controlled inline heater system (inline heater SH-27B, controller TC-324C, cable with thermistor TA-29, Warner instruments) to maintain the temperature in the perfusion chamber. A diaphragm was used to reduce the diameter of the excitation spot so that during imaging, only one cell at the center of the FOV was illuminated. To maximize photon collection, an oil immersion 40× NA 1.3 objective (CFI Plan Fluor DIC H/N2, Nikon Instruments) was used. A MATLAB (version R2024a, MathWorks) routine was used to control the PMT bias voltage and record the output voltage using the data acquisition and output boards. Data were collected at 80 kHz. The output voltage from the PMT was analyzed by a custom routine written in MATLAB to obtain the fluorescence signal for each cell. The raw data was first downsampled to 20 kHz.

We corrected the trace for photobleaching by dividing each trace by the best-fit three-term exponential (when the cell was held at −70 mV). The traces from the three replicate voltage steps were averaged. The corrected averaged signal was cropped from 0.1-s before the estimated depolarization or the repolarization onset to 1-s after the estimated depolarization or repolarization onset. The resulting kinetics traces were fitted with either a single-exponential,

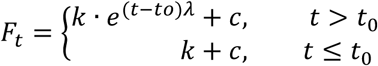

a dual-exponential model,

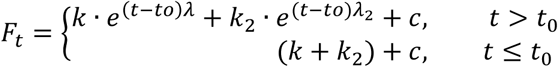

or a triple-exponential model,

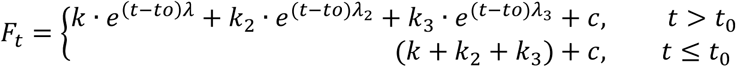

where *t* is the independent variable, time, and *F* is the dependent variable, fluorescence response. The coefficient *c* describes the mean plateau fluorescence, *k*, *k*_2_, and *k*_3_describe the relative ratio of each exponential component, *λ*, *λ*_2_, and *λ*_3_describe the negative inverse of the time constant(s), and *t*_0_is an offset indicating the exact event onset timing. For the −70 to +30 mV step, ASAP4e and FORCE1s were best fit by a triple exponential, where *λ*_3_ describes a slow photobleaching component not reported in Supplementary Table 1. For the +30 to −70 mV step, JEDI-2P and FORCE1s were best fit by a double and triple exponential, respectively, where *λ*_2_and *λ*_3_ describe a slow photobleaching component not reported in Supplementary Table 1.

#### Fluorescence lifetime

HEK293A Kir2.1 cells were cultured under standard conditions and seeded at 60–80% confluency onto 24-well glass-bottom plates (Cellvis, Mountain View, CA, USA). Cells were transfected with genetically encoded voltage indicator (GEVI) constructs (JEDI-2P, FORCE1s, and FORCE1f) using JetPrime reagent (Polyplus-transfection SA, Illkirch, France) according to the manufacturer’s instructions. For each well, the transfection mixture consisted of 50 µL JetPrime buffer, 0.5 µg GEVI plasmid DNA, and 1 µL JetPrime reagent, and cells were maintained for 48 h post-transfection prior to imaging.

Fluorescence lifetime imaging (FLIM) was carried out using a Nikon A1 laser-scanning confocal microscope (Nikon Instruments Inc., Tokyo, Japan) equipped with a PicoQuant LSM upgrade kit (PicoQuant GmbH, Berlin, Germany). Excitation was provided by a 488 nm diode laser (350 mW), and fluorescence was collected through a Nikon 20×Plan-Apo 0.75 NA objective. To select for GEVI fluorescence, a 520/35 nm bandpass filter was positioned in front of the single-photon avalanche diode (SPAD) detector used for FLIM acquisition. Data acquisition and lifetime analysis were performed using Nikon Elements software (version 5.21.03, Nikon Instruments Inc.) in conjunction with SymPhoTime64 software (version 2.7.5615, PicoQuant GmbH). Cells were imaged in extracellular solutions containing either 5 mM KCl, corresponding to a hyperpolarized membrane potential of approximately −90 mV, or 100 mM KCl, corresponding to a depolarized membrane potential of approximately 0 mV. For each condition, four independent fields of view were recorded, each containing 30–40 cells, and fluorescence lifetime values were quantified using SymPhoTime64 software.

#### Confocal imaging in dissociated neurons

E18 mouse hippocampal neurons were purchased from Transnetyx tissue by BrainBits (SKU SDEDHP, Transnetyx, Inc.) and came in Hibernate® EB complete Media as single-cell suspensions. Before seeding the hippocampal neurons, a 24-well glass-bottom plate (P24-1.5H-N, Cellvis) was coated with Neuron Coating Solution (027-05, Sigma-Aldrich) at 37 °C overnight and then washed three times with PBS. Neurons were seeded at 100,000 cells/mL in 500 µL glutamate-supplemented NbActiv1 medium (SKU NB1 + GLU, Transnetyx Inc.) per well of a 24-well plate, following the manufacturer’s protocol. The plating day was considered as DIV 0. Half of the media was henceforth replaced with fresh, pre-equilibrated NbActiv4 medium (SKU NB4, Transnetyx Inc.) every 3–4 days. AAV transduction was conducted on DIV 7 with 1 µL hSyn-FORCE1s in AAV2/1 for each well. After 2 hours incubating with AAV, the medium was changed.

Ten days after transfection (DIV 17), the attached neurons were washed twice with the imaging solution 2 before adding a final 500 µL per well as the imaging solution. Laser-scanning confocal images were obtained using a high-speed confocal microscope (LSM880 with Airyscan, Zeiss) driven by the Zen software (version 2.3 SP1 FP3 black edition, Zeiss). The microscope was equipped with a 40× 1.1 NA water immersion objective (LD C-Apochromat Korr M27, Zeiss), a 488-nm argon laser (LGK7812, Lasos) set to 5% power (∼10 µW) and a per-pixel dwell time of 3.02 μs. Emission light was filtered using a multipass beamsplitter (MBS 488/561/633, Zeiss) and acquired with a 32-channel GaAsP detector (Airyscan, Zeiss) with a detector gain of 800, and a 1.92 Airy unit pinhole size. Images were acquired at a resolution of 0.04 µm/pixel and an X-Y dimension of 5528 × 5528 pixels. Z stacks were stepped at 4.439 µm between images. To increase the signal-to-noise ratio, 2 scans were performed and averaged for each image. Airyscan processing was applied to the images via Zen software (version 2.3, blue edition, Zeiss) to increase the resolution. In Figure 1K, the overall image and dendrite zoom-in are from the maximum projection of the z-stack with 25 images, whereas the soma image is a single slice. Scale bars, 20 μm.

### Simultaneous electrophysiology and resonant-scanning two-photon voltage imaging in the mouse cortex

#### Animal handling

All procedures were carried out in accordance with the ethical guidelines of the National Institutes of Health and were approved by the Institutional Animal Care and Use Committee (IACUC) of Baylor College of Medicine. Mouse strains were sourced from our breeding colony. Mice were housed under standard conditions (12-h light/dark cycles, light on at 6 a.m., with water and food ad libitum)

#### Cranial windows, headbars, and viral injections

Cranial window surgeries were performed on wild-type C57BL/6J (RRID:IMSR_JAX:000664), VIP-Cre (RRID:IMSR_JAX:010908, or Sim1-Cre (MMRRC:031742-UCD) or Tlx3-Cre (RRID: MMRRC_041158-UCD) mice (7-16 weeks old). The VIP-Cre, Sim1-Cre, and Tlx3-Cre mice were used in experiments that did not require Cre-recombinant expression due to limited availability of wild-type mice. In total three male mice and two female mice were used in this study. Anesthesia was induced with 3% isoflurane and maintained at 1.5-2.0% throughout the procedure. Meloxicam (5mg/kg) was subcutaneously injected at the start of surgery for analgesia. Mice were positioned in a stereotaxic frame (Kopf Instruments or Neurostar Drill and Injection Robot), and body temperature was maintained at 37℃ using a homeothermic blanket (Somnosuite with RightTempModule, Kent Scientific). The scalp was shaved, and bupivacaine (0.05 cc, 0.5%, Marcaine) was injected subcutaneously at the incision site. After 10-20 minutes, a ∼1cm2 region of skin was removed above the skull. The exposed fascia was removed, and the wound edges were sealed with a layer of surgical adhesive (VetBond, 3M). A 13mm stainless steel washer (Seastrom) attached to a custom, removable headbar was implanted on the skull with dental cement (CCB Metabond). The washer was centered approximately 2.7 mm lateral to the midline and 1.5 mm anterior to the lambdoid suture, allowing optical access to visual cortex. After the cement hardened, the mice were removed from the stereotaxic head holder and fixed to a small stage while maintaining anesthesia. A 4 mm-diameter craniotomy was drilled at the center of the washer with a HP1/2 burr (Meisinger HM1-005-HP). The exposed cortex was rinsed and submerged in artificial cerebrospinal fluid (ACSF; 125 mM NaCl, 5 mM KCl, 10 mM Glucose, 10 mM HEPES, 2 mM CaCl2, 2 mM MgSO4). After the craniotomy was completed, virus was injected into the visual cortex via a nano-injection pump (WPI or Neurostar Drill and Injection Robot) at a rate of 5 nL/sec. The pipette was removed approximately 1.5 minutes after the injection was complete. For two-photon guided patching experiments, dura was left intact to reduce swelling. A 4mm glass coverslip (Warner Instruments) was sealed to the skull using cyanoacrylate glue (VetBond). Super glue (Loctite) was used to secure the window edges. All mice were singly housed and allowed to recover for two weeks post-operatively to allow for expression of the virus.

FORCE1s-Kv expression was transduced by injecting AAV-hsyn-FORCE1s-(GSS)_3_-Kv (titer:1.2E13 GC/mL) in WT mice (C57/B6). For some injections, the virus was diluted 1:10 with ACSF (final titer: 1.2E12 GC/mL). WT mice were injected with total volumes ranging from 350-700 nL of virus at approximately 350-450 µm depth from the surface of the cortex.

Traces shown in Figure 2C-G are from a whole-cell recording. Of the 11 cells analyzed for precision-recall and F1-score comparison (Figure 2H-I), seven were whole-cell and four juxtacellular recordings. Gigaseals were allowed to stabilize for 3-5 minutes before break-in. The mouse was kept anesthetized using 1-2% isoflurane, and its body temperature was maintained at 37°C using a homeothermic blanket. In vivo two-photon guided patching experiments of the 11 neurons were obtained from 3 male and 1 female mice.

Following confirmation of viral expression, the initial cranial window was replaced, with a custom coverslip that had a pre-drilled ∼0.5 mm hole (diamond-tipped burr, Coltene/Whaledent). Window replacement was performed at least two weeks post-operatively to confirm expression and optimize the placement of the pre-drilled coverslip. The pre-drilled coverslip provided access for patch pipettes to enter the brain while still maintaining stability for imaging, and was positioned to enable targeting of neurons expressing FORCE1s-Kv. To prevent leakage of glue onto the brain, the hole was either sealed with a thin layer of Kwik-Cast or temporarily covered with ACSF or Gelfoam (Pfizer Hospital 00009-0353-01). After the window was sealed, ACSF was placed on the window surface to act as external solution and prevent the brain from drying out.

#### Electrophysiological recordings

Patch pipettes were pulled from borosilicate glass (Sutter Instrument; 1.5 mm outer diameter, 0.86 mm inner diameter) to a resistance of 6–11 MΩ and filled with standard internal solution containing: 121 mM K-gluconate, 4 mM KCl, 10 mM HEPES, 10 mM Na₂-phosphocreatine, 4 mM Mg-ATP, 0.3 mM Na-GTP, and 13 mM biocytin (pH 7.25, osmolarity adjusted to match the extracellular solution). Alexa Fluor 568 (20 μM) was added for pipette visualization. Pipettes were advanced under two-photon guidance using high positive pressure (∼400 mbar) to penetrate the dura and low pressure (20-40 mbar) to approach target cells. Pressure was controlled via a manometer (Fisher Scientific 06-664-19) and a custom-built manifold that allowed rapid switching between high and low pressures, minimizing the ejection of internal solution. Recordings were acquired in current-clamp mode. Liquid junction potentials were not corrected. When giga-ohm seals were not achieved, juxtacellular recordings were obtained. In some instances, current was injected to evoke action potentials. Patch-clamp recordings were performed using a Scientifica PatchStar micromanipulator and an Axon Instruments CV-7B headstage. Signals were amplified with a MultiClamp 700B amplifier (Axon CNS Molecular Devices) and digitized at 10kHz using a National Instruments DAQ system (BNC-2090A interface) and recorded through a custom program written in LabVIEW 2016. A Quest Scientific HumBug noise eliminator was used in most recordings to suppress line noise.

#### Voltage imaging instrumentation for combined imaging and patching

All scans were recorded using a two-photon resonant-scanning system (ThorLabs) with excitation provided by a tunable Ti:sapphire femtosecond laser (Coherent; Chameleon Vision). The frame rate for all these experiments was 440 Hz and the excitation wavelength was 920nm. A 0.8 NA 16x objective (Nikon; N16XLWD-PF) was used for imaging. Output power was measured post-objective using ThorLabs PM100D meter and ranged from 34 to 105 mW.

#### Data exclusion criteria

Cells were excluded from the analysis if any of the following conditions were met: no discernible spiking activity was observed in the fluorescence trace; the fluorescence and electrophysiological traces were not clearly correlated, indicating that the imaged and patched cells were not the same; tissue damage occurred due to high laser power; or substantial Z-axis movement was detected during imaging. Patch stability can decline over time, and eventual loss of the patch or a deterioration of the cell’s health is expected. Recordings were therefore manually truncated to exclude periods corresponding to patch loss or cellular deterioration. Segments showing severe photobleaching were also manually truncated and omitted from the analysis. The remaining usable portions included in the analysis had a mean duration of 413.7 ± 208.1 s.

#### Data pre-processing

Following each two-photon imaging experiment, the scans were corrected for motion as described in the voltage imaging data preprocessing section below. For each recording, we manually segmented cells within the FOV. To increase SNR, the correlation between each pixels’ fluorescence and the simultaneously recorded patch-clamp voltage trace was computed. Then, the top-correlated pixels within a user-defined bounding box around the patched cell were averaged to obtain a single fluorescence trace per cell. Fluorescence traces were normalized as described in the following section.

For Figure 2C-D, F_0_ is defined as the bottom 5th percentile of the entire trace.

#### Optical spike inference

For Figures 2 C, E, and H-I, spikes were detected in the optical trace using an adaptive threshold modeled after Caiman Volpy’s adaptive threshold.^35^ We decided to use an adaptive threshold method to avoid manually setting thresholds specific to each trace to detect spikes (since this would not be possible in a normal experiment without ground truth data). Fluorescence traces were first high-pass filtered at 20Hz and then passed through the adaptive threshold function, which selects a threshold based on the statistical distribution of local peak amplitudes. After filtering, a first-pass peak detection was run to identify local maxima. To compute the detection threshold, we estimated the probability density of all peak amplitudes via a Gaussian kernel density estimator (KDE). The empirical distribution of the noise was modeled by reflecting the KDE around the median amplitude, assuming the noise is symmetric. We computed tail integrals over the original/true KDE (*F_max_*) and reflected/noise KDE (*F_noise_*) distributions at a grid of amplitude x and defined a score function:

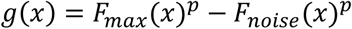

where *p* ∈ (0,1] is a stringency parameter (set to 0.1 in our analysis). The optimal threshold, *x*, was selected at the value that maximized *g(x)*. This threshold is then applied, and peaks were detected on points above the threshold.

For the precision-recall comparisons between JEDI-2P and FORCE1s loose patch recordings (Figure 2H), the adaptive threshold was scaled across a range of multiplicative factors. Because fluorescence amplitude distributions differed across recordings and indicators, the range of scale factors explored was adjusted per recording to ensure adequate sampling of the full precision-recall curve. The adaptive threshold computation itself (including filtering, KDE estimation, and *p*=0.1) was identical across recordings.

JEDI-2P recordings were obtained from the publicly released dataset accompanying the original JEDI-2P publication and analyzed using the same spike-detection and precision-recall workflow as for the FORCE1s recordings. The shared JEDI-2P dataset was provided as ΔF/F_0_ traces and was high-pass filtered at 20 Hz before spike detection. In contrast, FORCE1s’ raw fluorescence traces were high-pass filtered at 20 Hz before thresholding.

#### Correspondence between subthreshold voltages and FORCE1s-Kv responses

To examine the relationship between membrane potential and optical signal across a range of subthreshold membrane activity, both the electrophysiological and fluorescence traces were low-pass filtered at 50Hz to remove high-frequency noise (Figure 2F). The fluorescence trace was temporally interpolated to match the electrophysiological recording sampling rate. The resulting paired traces were then binned into a 150×150 two-dimensional histogram representing the probability distribution of ΔF/F_0_ values as a function of membrane voltage. The resulting bin widths were approximately 0.22mV (x-axis) and 0.0038 ΔF/F_0_ (y-axis) computed from the mean spacing between consecutive histogram edges. A linear regression was applied to quantify the relationship between voltage and fluorescence.

#### Precision-recall curves

Spikes were detected in both electrophysiological and optical traces. Singlets were isolated based on electrophysiological data, and spike times (10 kHz) were mapped to the nearest imaging frame center (440 Hz). Regions containing doublets or bursts were excluded. Each detected event was classified as a true positive (TP), false negative (FN), or false positive (FP). A TP was defined as an electrophysiological spike with a corresponding fluorescence spike within the same or subsequent two frames; an FN as an electrophysiological spike without a fluorescence counterpart; and an FP as a fluorescence spike without an electrophysiological match. Precision and recall were computed as

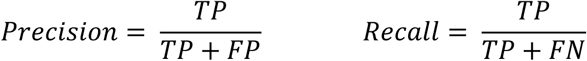

and F1 scores as

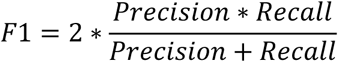

Precision-recall curves were generated across scaled thresholds to identify optimal parameters per cell, as well as a single optimal threshold that could be applied uniformly across the 11 cells (Figure S2.1A). The adaptive threshold was scaled to 0.1-2.2. For each recording, precision, recall, and F1 were computed at each scale factor.

For two of the JEDI-2P cells, multiple recordings were obtained. In these cases, the recordings were analyzed independently, and TP, FP, and FN counts were summed across recordings at each scale factor before computing precision, recall, and F1. This resulted in a single precision-recall curve per cell.

Within each curve, the scale factor that produced the highest F1 score was defined as the best threshold for that cell (Figure 2I, S2.1B). To select a single scaled adaptive threshold across the population of FORCE1s cells, we selected the scale factor that maximized the overall F1 score across all 11 cells (“Same thresholds all cells” in Figure S2.1B)

#### Spike-triggered averages

To align optical and electrical signals at the spike times, electrophysiological spike times (10kHz) were mapped to the nearest frame center (440 Hz).

Spike interpolation for baseline estimation: To prevent spike transients from biasing F_0_, a spike-removed trace was generated by linearly interpolating across detected electrophysiological spike time points.

Per-event ΔF/F_0_ baseline for STA: For each spike, a fluorescence snippet was generated for all regions of identified singlet spikes in the voltage trace. The baseline for the snippet was the mean of the last 20 frames (∼40ms) of the spike-interpolated fluorescence trace immediately before the spike, while excluding the final 5 frames just before the spike peak to avoid onset contamination. The snippet was then converted to ΔF/F_0_ using the per-event baseline and averaged across events to obtain the STA (SEM computed across snippets). Voltage snippets were aligned in time using patch timestamps, and a pre-spike mean from the spike-interpolated voltage was subtracted to express STA relative to its local baseline.

##### Resonant-scanning two-photon voltage imaging in the mouse cortex

#### Animal handling G surgeries

As in the previous section, except that dura was removed while avoiding damage to improve imaging quality. Cranial windows were implanted in wild-type C57BL/6J (RRID:IMSR_JAX:000664) or Tlx-Cre (RRID: MMRRC_041158-UCD) mice (7-16 weeks old). Four male mice and two female mice were used in these studies.

#### Voltage imaging instrumentation

Single-cell and multi-cell imaging scans shown in Figure 2K-O were acquired on a ThorLabs Bergamo resonant microscope with laser excitation provided by a tunable Ti:sapphire femtosecond laser (Coherent; Chameleon Discovery) tuned to 920 nm or a ThorLabs 2P-RAM mesoscope with a 920nm femtosecond laser (Toptica). The laser was delivered to the 2P-RAM breadboard via a hollow-core fiber, bypassing the standard precompensation path for the 2P-RAM scopeInstead, precompensation was applied internally in the laser via software. A dichroic mirror split emission light into two channels: the green channel used a 525/50 nm filter, and the red channel used a 625/90 nm filter, before being collected by two photomultiplier tubes. ScanImage (version 2018 Vidrio) was used to control the microscope and acquire imaging data. Imaging data on the Thorlabs Bergamo microscope was acquired with either a 16X 0.8 NA objective (Nikon; N16XLWD-PF) or a 25X 1.1 NA objective (CFI75 Apochromat 25XC W, Nikon Instruments). Resonant two-photon voltage and acetylcholine multiplex imaging scans were acquired on the previously described Thorlabs Bergamo microscope with laser excitation provided by a tunable Ti:sapphire femtosecond laser (Coherent; Chameleon Discovery) tuned to 1020 nm. A 16x 0.8 NA (Nikon; N16XLWD-PF) was used for imaging.

#### Viral injections

FORCE1s-Kv shown in Fig. 2K-O, Fig. 5A, 5B, 5G was transduced by injecting WT or Tlx-Cre mice with AAV-hsyn-FORCE1s-(GSS)_3_-Kv (titer:1.20E13 GC/mL), AAV-EF1α-DIO-FORCE1s-WPRE (titer: 1.05E13 GC/mL), or AAV-FORCE1s-TSER-Kv-WPRE (titer:7.137E12 GC/mL)). Cre-dependent FORCE constructs injected in WT mice were co-injected with AAV1-CaMKII-0.4.Cre.SV40 (Addgene viral prep # 105558-AAV; 1.1E13 GC/mL) in a 1:1-10 ratio. A total volume of 350-700nL was injected approximately 350 and 500 µm depth from the cortical surface.

For multiplexing experiments shown in 5G-J, FORCE1s-Kv and rACh1h expression was transduced in WT mice by injecting each construct in different sites in an overlapping manner. 500-700 nL of AAV-hsyn-FORCE1s-(GSS)_3_-Kv virus (titer:1.20E13 GC/mL) was injected 300-500 um away from an adjacent injection of 500-700 nL of a red acetylcholine GRAB sensor (AAV-hsyn-rACh1h) (titer: 1.45E13 GC/mL; Y.Li, Peking University; DOI:10.1101/2024.12.22.627112).

#### Sampling

Single-cell imaging data shown in Fig. 2K-O were obtained from 4 single-neuron recordings from 2 male and 2 female mice and were performed at depths ranging from 53 to 468 µm, with post-objective powers ranging from 20 to 55 mW, as measured using a ThorLabs PM100D meter. Imaging FOVs of data shown in Fig. 2K-O were composed of 24-32 lines with a resolution of 1-1.25 px/µm. Multi-cell imaging data shown in Fig. 5A include 6 neurons imaged simultaneously at 400 Hz at a depth of 263 µm with 35 mW of power as measured by a ThorLabs PM100D meter. The imaging FOV spanned 450 pixels in width by 194 pixels in height at 0.4 px/µm resolution.

The functional data shown in Fig 5C-F were acquired at depths of 263-273 µm and frame rates of 108-396 Hz. The imaging FOVs shown in Fig. 5C spanned 200×314 px and 250×392 µm in width and height, respectively. The data shown in Fig. 5E comprises 37 neurons recorded across two scans acquired from one mouse. Data shown in 5F is composed of 19 neurons recorded in a single scan acquired from one mouse. Imaging power was 35 mW, as measured at the output of the objective with a ThorLabs PM100D meter.

#### Behavioral recordings

Pupil and running activity were obtained concurrently with neuronal recordings across all mice. Treadmill activity was tracked via a rotary optical encoder (Accu-Coder 15T-01SF-2000NV1ROC-F03-S1) coupled to a freely moving circular treadmill and sampled at 100 Hz. Periods where the velocity exceeded 1 cm/s for at least 1 second were considered running periods.

Pupil recordings were acquired using a camera (Genie Nano C1920M, Teledyne Dalsa) with a 1” hot mirror (FM02 Thorlabs) at 20 fps. The hot mirror was positioned between the mouse’s left eye and the monitor to prevent obstruction of the monitor while visual stimulation was presented. Single-cell and multi-cell spontaneous activity scans and were performed without any visual stimulation. During these spontaneous activity scans, we used the monitor’s backlight or a UV LED to keep the pupil at a baseline level, allowing us to observe fluctuations in size. Pixelwise orientation tuning scans utilized a global directional parametric stimulus (voltage_monet) described at the end of this section. Following pupil imaging, a DeepLabCut model (version 2.0.5) identified and tracked pupil edges from which pupil diameter was computed and used to track the location and size of the pupil. These measurements are transformed into physical units using a conversion factor based on the standard size of the mouse eye as previously described^36^. Pupil frames with poor detail where the pupil was not tracked by DeepLabCut were marked with NaNs. Pupil periods where the mean dilation exceeded 1 second and constriction speed was greater than 0.2 mm/s were treated as unique pupil periods for analysis. Pupil periods that were composed of 25% or more NaNs were excluded and not analyzed. To focus our analysis on state changes during quiet wakefulness, we excluded pupil periods that occurred 3 seconds before, after, and during running. Custom MATLAB (version 2023b) and LabVIEW (version 2016, National Instruments) code was used to control the acquisition and synchronization of imaging and behavior data.

#### Visual stimuli

For pixelwise-tuning scans, mice were presented with Gaussian noise with coherent orientation and motion (monet_voltage) as described previously^37^. The stimulus has 16 directions of motion randomly interleaved, each repeated 100 times. Each motion presentation period lasted 0.5 seconds. The stimulus lasted approximately 14 minutes.

#### Data exclusion criteria

Recordings were excluded from the analysis if any of the following conditions were met: no discernible spiking activity was observed in the fluorescence trace; tissue damage occurred due to high laser power; or substantial Z-axis movement was detected during imaging.

#### Data preprocessing

Voltage imaging data was preprocessed by using a pipeline implemented in Python (3.6.9). Imaging data were motion-corrected in the horizontal plane for motion shifts in x- and y-using a two-step algorithm. First, large motion shifts were corrected by computing a global average template across the entire scan and then computing the phase correlation between the global average and individual frames. Secondly, smaller motion shifts were corrected by computing a local template composed of different-sized windows averaged (3 to 30 seconds, depending on which performed best) and computing the phase correlation between the local template and individual frames. Scans with average x-y motion exceeding 10 µms, sudden artifacts, or loss of fluorescence due to water loss or external light contamination were excluded from analysis. Following motion correction, individual soma were manually segmented from the motion-corrected summary image. Raw fluorescence traces were then extracted of the manual segmented masks from the motion corrected imaging data.

Subsequently, extracted single and multi-cell neuronal traces were normalized to ΔF/F_0_ as described below:

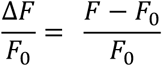

For Fig. 2L-N, and 5B-C, F_o_ was computed using the bottom 5th percentile of fluorescence values over 30-s windows. For Fig. 5A, no baseline correction was performed.

#### Characterization of single-cell and multi-cell FORCE1s activity

To characterize the photostability of FORCE1s spiking shown (Figure 2L-O), spikes were detected by passing a threshold parameter to the find peaks function in the scipy.signal Python package (version 1.2.3). To determine the thresholds to use, we computed the 99.8th percentile of each normalized trace with the percentile function within Numpy Python package (version 1.19.5). Singlets were identified by finding isolated detected spikes that had at least 100 ms before and after another detected spiking event. Spike triggered average shown in Figure 2N is of detected singlets (n=1207) in the detrended trace shown in 2K. Trace samples 500 ms before and after detected singlets were obtained from the ΔF/F_0_ trace. Trace samples were then normalized by subtracting each trace sample by its mean from −500 ms to 0 ms. Normalized trace samples were subsequently averaged to obtain the spike triggered average. For computing the SNR of detected singlets (Figure 2O, n=4 neurons from 3 mice), 200 ms periods of the raw trace, centered on detected singlets, were normalized by subtracting each period by the mean of the-75 to −25 ms period preceding each singlet. We defined signal as the peak of the normalized FORCE1s singlet trace and the noise as the standard deviation of the period −75 to −25 ms preceding the peak of the singlet. Subsequently, we computed the SNR for each detected singlet and averaged the values for minutes 1-5 and 25-30 for single-cell scans that spanned at least 30 minutes.

Additionally, we computed the normalized rolling mean spike rate for each trace across the duration of the scans (Figure 2M, n=4 neurons from 3 mice). Mean spike rate was computed using a 1-minute rolling window with 250 ms steps. The mean spike rate for each trace was normalized to the mean spike rate of the first minute.

#### Pixel-wise analysis of orientation tuning

For pixel-wise analysis orientation tuning analysis (Figure 5B) motion correction was performed, and the scans were normalized in two stages. First, the data was detrended by subtracting a 0.1 Hz low pass filtered version of the scans and then dividing by it. Each pixel timeseries was centered on their mean and then dividing by the root-sum-of-squares of each pixel’s temporal component. To map each pixel’s preferred direction, a linear regression model was used. For each unique direction, a regressor *R(t)* was designed:

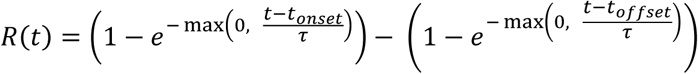

where *t* is the time of the frame, *t_onset_* is the start time of the trial, *t_offset_* is the end time of the trial, and τ is a time constant, set to 0.7. The regression coefficients, which represent the magnitude of the response to each direction, were calculated by multiplying the detrended scan with the regressor matrix. For each pixel, a single complex number was computed by projecting the N regression coefficients onto a complex-valued basis vector. The basis vector was constructed using the expression *e*^2i^*^θ^*,where θ is the direction in radians, and was scaled by a factor of *1/√(N/2)*, where N is the number of unique directions. The orientation map was calculated from the phase angle of this complex number, with each pixel representing the preferred orientation of motion that elicited the strongest response. The amplitude map was calculated from the magnitude of the complex number, which quantifies the strength of the orientation tuning. The orientation map was further color-coded to visualize the results, with hue representing the preferred orientation and brightness and saturation representing the tuning strength from the amplitude map. To produce orientation tuning curves, the mean ΔF/F_0_ for each orientation of motion was computed.

#### Correlation between voltage and acetylcholine indicators’ responses

To compute changes in low-frequency activity correlated with changes rACh levels (Figure 5E), we binned changes in FORCE1s phase to changes in phase of rACh (n=37 neuron across two scans from one mouse). Previous publications have shown a high-correlation between brain state changes in low-frequency activity using electrophysiology and external behavior measures such as locomotory and pupil activity but have yet to establish whether cortical neuromodulator levels measured via optical sensors also exhibit brain state transitions reflected in pupil activity.^9,20,36,38–40^

To compute rACh phase binning of low-frequency activity, normalized FORCE1s traces were bandpass filtered using a zero-phase 2-10 Hz Hamming bandpass filter. A Hilbert transform was then applied to the band-pass filtered FORCE1s trace using the scipy.signal.hilbert function (version 1.2.3). The amplitude of the Hilbert transform was subsequently filtered between 0.1 and 1 Hz to match the timescales on which distinct dilating and constricting pupil periods occur. Additionally, the raw rACh trace was filtered between 0.1-1 Hz with a Hamming Bandpass filter, prior to computing its phase using the scipy.signal.hilbert trace. We then iterated through all non-running pupil periods and binned the filtered FORCE1s Hilbert transform amplitude by the phase of the filtered rACh (65 phase bins from -pi to pi). The resulting FORCE1s values binned by rACh phase were averaged, and error bars computed using the SEM. Finally, the phase-binned signals were then smoothed with a rolling average window of 3 bins.

Cross-correlations were calculated by first band-pass filtering, detrended rACh1h and FORCE1s fluorescence traces between 0.1 and 2 Hz (Figure 5J, n=19 neurons from a scan in one mouse). Filtered fluorescence traces were normalized by subtracting the mean and dividing by the standard deviation. Cross-correlations were then computed on the normalized traces. The lag at peak cross-correlation was determined within a three-second window centered on zero lag.

### AOD-based recordings using ULoVE or diagonally scanned imaging in the mouse cortex

All protocols adhered to the guidelines of the French National Ethics Committee for Sciences and Health, as outlined in the report on Ethical Principles for Animal Experimentation, in agreement with the European Community Directive 86/609/EEC, under agreement #29791.

#### Animal handling, viral injections, and surgeries

Male wild-type C57BL/6J adult mice (>P40 - body weight 20–24 g, n=4 for JEDI2P-Kv and n=6 for FORCE1s-Kv) were housed in standard conditions (12-hour light/dark cycles, light on at 7 a.m., with water and food *ad libitum*). As a preoperative analgesic, buprenorphine, 0.1 mg/kg, was used, and Ketamine-Xylazine was used as an anesthetic (Centravet). The expression of JEDI2P-Kv was from capsid type1 AAV viruses produced^41^ in-house at IBENS from Addgene clone #179459 using a plasmid ratio (µg) of transgene:capsid:helper: 1:1.6:2. For FORCE1s-Kv expression (except for the results in Fig 2D), virus was packaged into type1 AAV by VectorBuilder from a construct in their Cre-dependent backbone (pAAV.EF1a.FlexON.FORCE1s-Kv). FORCE1s.-Kv expression in Figure 2D was from clone 4934_pAAV.EF1a-DIO-FORCE1s-codonOptomized-Kv, and was packaged as described for JEDI2P-Kv. To provide GEVI soma localization, JEDI2P-Kv and FORCE1s-Kv were tagged with the PRC motif of Kv2.1, which was only codon optimized for JEDI2P-Kv. The Kv motif was attached to the C-terminus of the GEVI with a GSSGSSGSS linker (JEDI2P-Kv) or for FORCE1s-Kv, without this linker but with intervening Golgi and ER export motifs of Kir2.1 (Gradinaru et al. 2010). All GEVI viruses were CRE-conditional enabling sparse GEVI expression by co-injecting a CRE-expressing virus (final titer concentrations of 4.1e12 GC/ml GEVI virus with 3e9 GC/ml of AAV2/1 hSyn.Cre.WPRE.hGH (Addgene, 105553)) into the neocortex and at a depth of 350µm. To obtain cell-type specific, of GEVI-expression in pyramidal neurons, AAV2/1CamKII 0.4.Cre.SV40 (Addgene, 105558) was co-injected at a final titer of 3.1e10 GC/ml while the GEVI viruses were injected at a titer of 3e12 GC/ml.

GEVI and Cre AAV were combined in a saline solution containing 0.001% of pluronic acid (ThermoFischer 24040032), 300 nl of which was injected at a flow rate of 100 nl/min into the neocortex at two injection sites more than one millimeter apart. A custom-designed aluminum head-plate was fixed on the skull with layers of dental cement (Metabond). A 5-mm diameter #1 coverslip was placed on top of the visual cortex and secured with dental cement (Tetric Evoflow). Mice were allowed to recover for at least 15 days before recording sessions and were housed in groups of at least 2 mice per cage. Recordings were done between 21 and 60 days post-surgery. Behavioral habituation was adopted, involving progressive handling by the experimenter with gradual increases in head fixation duration^42^. Mice were handled before recording sessions to minimize restraint-associated stress, and experiments were conducted during the light cycle.

#### ULoVE voltage optical recording and experimental design

Recording sessions lasted 1-3 hours and were conducted while mice behaved spontaneously on top of an unconstrained running wheel in the dark^2^. Recordings were performed using an acousto-optic deflector (AOD)-based random-access multiphoton system (Karthala System) based on a previously described design^3^. The excitation was provided by a femtosecond laser (InSight X3, Spectra Physics) mode-locked at 920 nm with a repetition rate of 80 MHz. A water-immersion objective (CFO Apo25XC W1300, 1.1 NA, 2 mm working distance, Nikon) was used for two-photon fluorescence excitation and epifluorescence light collection. This experiment was done at 920 nm because the peak 2P excitation wavelength for the JEDI3 sensors had not yet been determined. The signal was passed through an IR blocking filter (TF1, Thorlabs), split into two channels using a 562 nm dichroic mirror (Semrock), and passed to two H12056P-40 photomultiplier tubes (Hamamatsu) operating in photon-counting mode. The 510/84 filtered green channel was used for collecting JEDI2P-Kv and FORCE1s-Kv signals. The laser power was set to deliver 15 mW post-objective and pre-sample, then adjusted for mono-exponential loss through tissue with a length constant of 170 µm. To locate neurons, before making ULoVE optical recordings, the AOD-based microscope was used in sequential point scanning mode to form an image. To produce the images in Fig. 3A,F, time series of 50 images (1 µs/pixel, 0.091 µm/pixel) were acquired and post hoc motion registered. Cell depth was measured from the brain surface.

The optimized ULoVE excitation pattern (Lombardini et al., in preparation) consists of a series of 9 points vertically aligned (evenly spaced over 15 µm), multiplexed twice horizontally with a 2 µm spacing. The 18 points were scanned diagonally (8 µm horizontally and 3 µm vertically) during a 50-µs acquisition time, to homogeneously fill an extended excitation volume that continually encompasses the cell plasma membrane. This latest strategy refines the axial profile of excitation used previously (Villette et al. 2019) thereby limiting signals outside the desired focal plane and reducing neuropil signal contamination, which improves SNR. Compared with the power used in the imaging mode, laser power was multiplied by 1.5 to account for the greater excitation volume. The applied power never exceeded 200 mW. Using two patterns per cell enabled a temporal resolution of 7142.9Hz, calculated as 50µs/volume plus 20µs of access time (multiplied by 2 because two volumes were used/acquisition). Recordings were stopped after 600 seconds (10 minutes). For recording, we selected neurons that were sufficiently bright to obtain a significant signal-to-noise ratio, yet did not display long-lasting depolarizing plateaus, which we believe indicate over-expression.

#### Signal analysis, spike extraction, and waveform analysis

For each cell recording, two ULoVE volumes were used as “regions/volumes” of interest; thus, the traces represent the sum of photons collected from the two volumes. The photon flux is the amount of photons per seconds. Because FORCE1s-Kv is a dim-to-bright GEVI, reliable recordings were found from cells that had rather low levels of baseline photon flux. This is in contrast to recordings of cells expressing the bright-to-dim GEVI, JEDI2P-Kv, where cells of brightness relatively higher on average than FORCE1s-Kv were found to produce reliable and robust recordings. Photobleaching was assessed by bi-exponential fitting and corrected by division of the raw trace by the normalized fit function. This analysis enabled the full trace to be corrected for the fast-photobleaching component. To characterize the second phase (from 60 seconds to the end of the recording), which could be more a consequence of a diffusion process described by a power law, the traces were low-pass filtered at 1 Hz. Traces were down-sampled 140 times and plotted on a log10-log10 scale, then fitted with a line, to determine the slope of the power law. The following steps were used to generate the %ΔF/F_0_ traces: the raw traces were normalized by the bi-exponential fit function (computed above), and the remaining low-frequency drift was removed using a zero-phase distortion filter (high pass: 0.5 Hz). The traces were then converted to %ΔF/F_0_, with the mean signal used as F_0_.

Spikes were detected using a custom-designed algorithm. First, each trace is high-pass filtered (using a second-order Butterworth filter with a lower limit of 40 Hz). Then we convert this trace with a delayed differential trace that is the difference between the “signal”, (defined as the trace’s mean over 2 ms) and the ‘baseline’ (defined as the trace’s mean over 3 ms and delayed by 1 ms). The rationale behind this is to define a baseline before the spike that does not encompass data points in the rising phase of the putative spikes. Choosing a baseline this way enables increasing the contrast both for isolated spikes and spikes within bursts. A subsequent step consists of enhancing the contrast by multiplying this delayed differential trace by the positive high frequency (250Hz cut off) content of that trace. Finally, this sharpened and highly contrasted trace is z-scored. Events above 20 std were considered as spikes. Lastly spike onsets are extracted as the local peak of the linear regression slope over a period of 3 ms on a 10 kHz linear interpolation trace.

To quantify spike amplitudes (Fig. 3B), an average spike waveform was extracted from isolated spikes. Single spikes that were separated by more than 50 ms were aligned to the spike onset and then averaged to produce the average spike waveform (Fig. S3.2A, green traces). Spike amplitude was then extracted from the peak value of the average spike waveform obtained only from isolated spikes measured from the onset point. FWHM corresponds to the extent in time at half maximum amplitude, taken from the average spike waveform after 20 kHz linear interpolation. Tau off corresponds to the time constant of an exponential fit on the repolarization phase of the spike. D’ is obtained as previously^4^. To get the spike SNR, the spike amplitude is divided by the shot noise, corresponding to the square root of the mean photon count of the trace.

To quantify subthreshold fluctuations, traces were filtered using a bidirectional Butterworth bandpass filter with a frequency range of 0.1 to 50 Hz. We restricted our analysis to time periods when the animal’s speed was below 1 cm/s. The cell’s low state corresponds to the 1^st^ percentile of that filtered trace. The cell’s high state corresponds to the median of the signals at the detected spike onsets. The subthreshold fluctuation is defined as the difference between the cell’s low state and its high state. To get the subthreshold SNR, this subthreshold fluctuation measure is divided by the shot noise (same as for spike above).

#### Downsampling of ULoVE recordings

To assess the impact of downsampling on spike amplitude detection, we analyzed extracellular waveforms originally sampled at 7.1 kHz. Waveforms were upsampled to 1 MHz and then downsampled to rates between 25 and 7000 Hz. For each sampling rate, we tested 20 temporal phases by systematically shifting the downsampling starting point. Peak amplitudes were normalized to the original waveform maximum. We quantified amplitude loss and variability (coefficient of variation) across neurons and phases for both JEDI and FORCE1s GEVI recordings.

#### Diagonally scanned image recording

AOD-based imaging by point-by-point scanning is inherently slow due to the 20 µs access time per pixel when controlling AODs at constant ultrasonic frequencies. To circumvent this limitation, a method based on controlling AODs with linear frequency chirps was implemented and termed “diagonally scanned image recording”. Applying a ramp of frequencies equally to the AODs that scan in the x and y directions produces a diffraction-limited PSF that is diagonally scanned (Lombardini, in preparation). The diagonal position and length are tunable, such that an ROI of any shape can be applied to the sample by a series of adjacent diagonal scans. Although the exposure time for each pixel along a diagonal is invariant once the scan commences, it can be set to as low as 0.1 µs before the scan begins.

Thus, we achieved imaging of 8 to 11 ROIs covering each a neuronal soma, and its close surrounding area (for post hoc motion registration), at a sampling rate above 200Hz using a pixel size of 0.909 µm and exposure time of 0.2 µs per pixel. To achieve a larger SNR, power was at most tripled compared to the reference power described above (45 mW at the surface). As scanning speed is rapid, and molecules are exposed only every 4-5ms, we observed no bleaching. To avoid computer memory limits during 1h long acquisitions, 61 repetitions (each lasting 1 minute and consisting of 15000 images) were acquired and saved in ‘raw’ format every minute with a saving time of <400ms between each. Movement correction was performed for each frame using cross-correlation to a reference image before traces were recorded. Traces are the summed photons from pixels that defined a given cell. To convert a trace in %ΔF/F, the trace has been subtracted from its mean and normalized by its mean then multiplied by 100.

### FACED2.0-based voltage imaging in the mouse cortex

#### Animal handling, viral injections, and surgeries

All animal experiments were performed in accordance with the National Institutes of Health guidelines for animal research. All procedures involving mice were approved by the Institutional Animal Care and Use Committee at the University of California, Berkeley.

Procedures for virus injection and cranial window implantation followed previously established methods^43^. Wild-type C57BL/6J or Rbp4Cre (+/-) mice were anesthetized with isoflurane (1-2% in oxygen) and provided buprenorphine (0.3 mg/kg, subcutaneous) for analgesia. Animals were stabilized in a stereotaxic frame (Model 1900, David Kopf Instruments), and a circular craniotomy of 3.5 mm was opened above the left visual cortex with the dura kept intact. A beveled glass pipette (15-20 µm tip, 45° angle), mounted on a hydraulic manipulator (MO10, Narishige) and backfilled with mineral oil, was used to inject pAAV2/9-hsyn-FORCE1s-(GSS)_3_-kv2.1 solution at 6-12 sites (20 nl per site) and a depth of 250 µm. Following injections, a single glass coverslip (No. 1.5, Fisher Scientific) was placed over the craniotomy and fixed with dental acrylic. A titanium head post was then secured to the skull using cyanoacrylate glue and dental acrylic. After a three-week recovery period, in vivo imaging was performed on awake, head-fixed mice.

#### Microscope and imaging parameters

The ultrafast two-photon fluorescence microscope was described previously^8,44^. Briefly, light from a 1035 nm fiber laser (Monaco 1035-40-40, 1 MHz repetition rate; Coherent) was focused by a cylindrical lens and directed onto a pair of mirrors with a small tilt angle. Through free-space angular-chirp-enhanced delay (FACED), each laser pulse was split into multiple pulses that were spatially separated and temporarily delayed, forming a 1D array of excitation foci along X axis at the objective focal plane. The FACED foci array was scanned by a galvo mirror along the Y axis to generate one FACED FOV of 80 µm × 500 µm at 1538.4 Hz using a 25× 1.05 NA objective (XLPLN25XWMP2; Olympus). To extend the FOV, multiple FACED FOVs were tiled, yielding a frame rate of 256.4 Hz across a 480 µm × 500 µm region. The tiled FACED FOVs were post-processed with Y-axis shifts to better align structures and X-axis stitching to remove overlapping regions. The final FOV size was reported for each figure. The post-objective excitation power was 185-198 mW. Imaging was performed in the left visual cortex of the mouse, at depths ranging from 90 µm to 110 µm.

#### Visual stimuli

A visual stimulation protocol was implemented in MATLAB (Mathworks) using the Psychophysics Toolbox^45^. The stimulation consisted of drifting gratings presented on a monitor. Each stimulation cycle began with a 0.5-second blank screen, followed by a 0.5-second presentation of a drifting grating. Grating direction varied from 0° to 315° in 45° increments. Each trial included all 8 directions and had a total imaging duration of 8.2 seconds.

#### Data analysis

Data were analyzed using custom MATLAB (Mathworks) programs. Time-streaming images were motion corrected using an iterative cross-correlation method^43^. ROIs were identified through a combination of automatic 2D cell segmentation (Cellpose 2.0^46^) and manual correction, and the average signal within each ROI was extracted as the raw trace.

ΔF/F was calculated by defining the baseline signal (F) as the mean fluorescence signal from 0.3 to 0.5 seconds during the blank period, with ΔF defined as the difference between the raw trace and F. For spike detection, the ΔF/F trace was detrended with a high-pass 3rd-order 15-Hz Butterworth filter, and peaks exceeding 3.5 times the sliding standard deviation (std) of the detrended ΔF/F were identified as valid spikes. To determine this spike-detection threshold, the ΔF/F trace was inverted, and the same analysis was applied. A threshold of 3.5 produced 1% false positives and was therefore used here (Figure S4.1B). Subthreshold activity was assessed by low-pass filtering the raw ΔF/F trace with a 3rd-order, 15-Hz Butterworth filter.

To assess whether a cell was visually responsive (VR) based on spiking activity, a one-tailed t-test was conducted to compare the firing rates during each of the 8 stimulation periods (0.5 to 1.3 seconds) with those during the blank period (0.3 to 0.5 seconds). To assess whether a cell was visually responsive based on their subthreshold activity, the one-tailed t-test was performed using the mean subthreshold ΔF/F values during the blank versus stimulation periods. An additional 0.3 seconds following the stimulation was included in this analysis to account for neuronal responses to the offset of grating stimulation. Cells with a p-value below 0.0063 (0.05/8, with multiple-comparison correction) were classified as VR.

For analyses evaluating how acquisition frequency affected spike detection, we temporally downsampled datasets by removing frames. All recordings—original and downsampled—were high-pass-filtered with a 15-Hz 3^rd^-order Butterworth filter. As above, ΔF/F was calculated by defining the baseline signal (F) as the mean fluorescence signal from 0.3 to 0.5 seconds during the blank period. Peaks exceeding 3.5 times the sliding standard deviation (std) of the ΔF/F traces were identified as valid spikes. ΔF/F was converted to Z-scores by dividing by the standard deviation of the signal from 0.3 to 0.5 seconds during the blank period. Spike Z-scores were computed by calculating the maximal Z-score for each spike identified in the ΔF/F traces. Per-cell spike counts were calculated by averaging across all trials.

### MINI2P-based cortical two-photon voltage imaging in freely moving animals

#### Animals handling

A total of 2 male wild-type (C57BL/6JBomTac) mice, aged 2-6 months, were used in the experiments, performed according to the Norwegian Animal Welfare Act and the European Convention for the Protection of Vertebrate Animals used for Experimental and Other Scientific Purposes, permit number 29894. Littermates were housed together in individual cages with 1-4 mice per cage, maintained on a regular diurnal lighting cycle (12:12 light:dark) with *ad libitum* access to food and water and nesting material for environmental enrichment. Mice were housed one per cage after surgery.

#### Surgeries and viral injections

To record from L2/3 V1 neurons, we followed the procedure described previously^24^. Briefly, a 3mm craniotomy was made over the left hemisphere (AP: −2.8 mm, ML: −2.5 mm), and AAV9-hSyn-FORCE1s-Kv (∼2 × 10¹¹–1 × 10¹² vg/ml) was injected at 4 locations (DV: 350 µm) separated at least 250µm from each other. Each injection consisted of 100 nl delivered using a nanoliter injector (Nanoinject III, Drummond Scientific) at a rate of 1 nl/s. Then, a 3mm-diameter coverglass was implanted.

#### MINI2P recordings

The MINI2P system used for imaging was as described previously ^24^, comparable to the Mini2P system available from ThorLabs. The laser source was a compact, single-wavelength femtosecond laser (Alcor-920-2, Spark Lasers, France: 920 nm, ∼100 fs, 40 MHz, 2.5 W). To achieve higher frame rates, a MEMS mirror (A7M10.2-1000AL, Mirrorcle, USA) running at a 6.4 kHz scan rate on the fast axis (x) was used. Also, small fields of view (FOVs) containing 1-3 neurons were imaged at resolutions of 32×24 to 128×24 pixels, enabling frame rates 414-427 Hz. Laser power was kept at 50-65 mW, measured under the objective.

Before the open-field task, voltage sensor expression was verified in head-fixed mice. Once a suitable FOV with neurons expressing the sensor was identified within a 300×300 µm area, mice were baseplated to attach the miniscope using a stitching adapter^24^. ScanImage software was used to control the MINI2P system, capture imaging data, and synchronize it with the animal tracking camera.

#### Open field task and animal tracking

Experiments started four weeks post-surgery, following full recovery. Mice were habituated to handling and the behavioral task for one week prior to the experiments. Two to three days before the experiments, they were acclimated to the miniscope’s weight by wearing a dummy version in their home cage. The task was performed in a dark room illuminated by a warm-light LED strip, synchronized with two-photon imaging to prevent light contamination. Neural activity was recorded as the mice explored an 80×80 cm black square box, with a single white A4 paper on one wall serving as a visual cue. Exploration was encouraged by periodically scattering cookie crumbs in the box. Each session consisted of a single 40-minute recording per mouse, with brief breaks (7–100 s; mean 40 s) every 10 minutes to detwist the miniscope fiber bundle when necessary and to save the data in smaller segments, which facilitates memory use during processing. We did not observe any noticeable change in the SNR or photobleaching speed due to the addition of these short breaks. Animal behavior was recorded using a monochrome NIR camera (acA2040-90umNIR, Basler) mounted above the box with a 16-mm lens (C11-1620-12M-P f16mm, Basler). The environment was illuminated with an 850 nm LED array, and an 850 nm bandpass filter (FBH850-40, Thorlabs) on the camera lens prevented interference from the 920 nm laser. The tracking video was recorded using Basler’s Pylon software in external mode. Synchronization with 2P imaging was achieved using 30 Hz TTL pulses (0-5V) generated by the vDAQ card.

#### Behavioral processing

The raw tracking videos were processed using DeepLabCut^47^, which provided the positions of the mouse’s body parts in each frame. The miniscope and four body parts were tracked: the left ear, right ear, body center, and tail base. The positions were filtered to include only data points with a likelihood > 0.7 and upsampled to match the 2P imaging frame rate using linear interpolation. To calculate the mouse’s speed in the arena, the Euclidean distance between consecutive positions of the tracked miniscope was divided by the inter-frame time interval. The resulting speed was smoothed using a moving average with a 0.5s kernel. The head direction was defined as 90 degrees counterclockwise relative to the vector from the left ear to the right ear.

#### Voltage signal analysis

All recordings were rigid motion-corrected with NoRMCorre^48^. The 40-minute recordings, composed of four 10-minute segments, were first concatenated before performing motion correction. Neuron ROIs were manually drawn from the mean image projection of the optical recordings, and raw fluorescence traces were extracted as the average pixel intensity within each ROI. To detect spikes, neuron Regions-Of-Interest (ROIs) binary masks were fed to Volpy pipeline^49^, which simultaneously inferred optimal pixel weights, spike timings, subthreshold signals, and ΔF/F trace.

To calculate the signal photobleaching of each neuron, raw fluorescence traces without spikes were generated by replacing the intensity values of the spike’s locations and surroundings (5ms before and 10ms after the spike) with the average fluorescence intensity within a local ±50ms time window. Then, baseline traces were estimated by low-pass filtering the spikeless traces at 1Hz. Photobleaching was estimated by normalizing the baseline trace to t = 0.

SNR was calculated similarly to previously described^50^. Briefly, the raw fluorescence signal was high-pass filtered at 0.25 Hz to remove possible slow intensity drifts, yielding a detrended trace. A subsequent high-pass filter at 20 Hz was then applied to remove subthreshold fluctuations, leaving primarily spike-related events and high-frequency noise. To estimate noise independently of spike activity, a local mean was computed over a ±1 s window at each time point, and any values exceeding this mean were replaced by the mean, thus isolating only downward deviations. The noise at each time point was defined as twice the standard deviation of these downward deviations calculated over a ±5 s window. The SNR trace was obtained by dividing the detrended signal by this dynamic, spike-independent noise measure. Finally, the SNR over time was determined by averaging the SNR values at spike locations within each one-minute interval.

#### Spatial tuning maps

Spatial tuning maps were generated to display the firing rates of neurons, normalized by occupancy. The value of each bin in the map was calculated as the ratio between the total number of spikes (predicted by VolPy) and the time spent in the bin (bin size: 2.5 × 2.5 cm²). The resulting maps were smoothed with a Gaussian filter (standard deviation: 3 cm). A bin was considered visited if the animal spent at least 0.1 s within it). To avoid confounds related to changes in behavioral state, all data corresponding to running speeds below 2.5 cm/s were excluded from the analysis.

Border score was calculated similarly as previously described^51^. In brief, firing fields were defined as contiguous regions with a minimum size of nine bins (56 cm²) and firing rates at least 30% of the session peak. For each wall, border coverage was computed as the fraction of valid wall-edge bins in direct contact with a firing field, and the maximum coverage across the four walls was taken as the coverage measure. The mean firing distance was then calculated as the firing rate–weighted average distance of field bins to the nearest wall. A border score was defined by comparing this distance with the maximum wall coverage, yielding values from −1 for fields in the center to 1 for fields aligned with a wall.

The significance of spatial tuning was evaluated using a shuffling procedure. The entire spike sequence of each neuron was circularly time-shifted along the animal’s trajectory by a random interval between 30 s and the session duration minus 30 s. This procedure was repeated 200 times to generate a null distribution of spatial tuning scores, ensuring that tuning was not due to chance. A neuron was classified as spatially tuned if its observed score exceeded the 95th percentile of the shuffle-derived distribution. For border cell analysis, neurons with a border score above 0.5 were considered border-tuned.

## Data and code availability

- The sequence of FORCE1s is available from GenBank [accession number to be provided prior to publication].
- Plasmids used for AAV packaging for *in vivo* voltage imaging are available from Addgene [Addgene IDs to be provided prior to publication].
- Processed data for all the main figures are deposited [final data will be uploaded to Zenodo or provided as a Supplementary file]
- Raw data for main and supplementary figures are deposited [final datasets to be uploaded to Zenodo prior to publication].
- Data that is too large to upload will be made available upon request.
- This paper does not report original code needed to analyze the data generated by this study.
- Any additional information required to reanalyze the data reported in this paper is available from the lead contact upon request.

## Acknowledgments

**St-Pierre lab.** We thank Zhuohe Liu and Anh Pham for assisting with GEVI screening. We also thank M. E. Dickson, J.M. Kirk, C.-W. Hsu and the Imaging & Vital Microscopy Core (Baylor College of Medicine, BCM) for training on the confocal microscope. The project described was supported in part by the Neuroconnectivity Core at Baylor College of Medicine, which is supported by IDDRC grant number P50103555 from the Eunice Kennedy Shriver National Institute of Child Health & Human Development. The content is solely the responsibility of the authors, and it does not necessarily represent the official views of the Eunice Kennedy Shriver National Institute of Child Health & Human Development or the National Institutes of Health. Fluorescence lifetime imaging microscopy was performed at the Center for Advanced Microscopy, a Nikon Center of Excellence, at McGovern Medical School, UTHealth Houston (RRID: SCR_025962). **Reimer lab.** We thank Cameron L. Smith for help analyzing orientation-tuning and acetylcholine experiments, and Y. Li (Peking U.) for providing AAVs expressing an Ach indicator. **Bourdieu lab**. We thank Caroline Mailhes-Hamon for helping to establish dense labeling and Bertrand Ducos for virus titration. **Grant support.** The project was supported by a Vivian L. Smith Endowed Professorship in Neuroscience (FSP); a Klingenstein-Simons Fellowship Award in Neuroscience (FSP); the McNair Medical Foundation (FSP); Welch Foundation grant Q-2016-20190330 and Q-2016-20220331 (FSP); NIH grants R01EB027145 (FSP), R01EB032854 (FSP), U01NS113294 (FSP, ML), U01NS118288 (FSP), U01NS133971 (FSP, LB), R01NS136027 (FSP, JR), R01NS146023 (FSP, JR), R01NS146078 (FSP, JR), U01NS118300 (NJ), U01NS137449 (NJ), Weill Neurohub (NJ), RF1NS128901 (JR), R34NS132045 (JR); NSF NeuroNex grant 1707359 (FSP), IdeasLab grant 1935265 (FSP), The Research Council of Norway FRIPRO grant ES737113 (WZ), Proof of Concept grant 355770 (WZ), Centre of Excellence grant; Center for Algorithms in the Cortex 332640 (WZ), and the National Infrastructure grant NORBRAIN IV (WZ), Agence Nationale pour la Recherche ANR-25-CE16-7612-01 (LB).

## Author contributions

**St-Pierre lab.** FSP conceived the project. FSP, AJM, MAL, and SY coordinated the project. AJM, MAL, and SY screened GEVIs. MS, AJM, and MAL optimized screening protocols or hardware. XD, HA, RB, and SL constructed mutagenesis libraries and viral vectors. GM, ZL, HL, BC, KLC, and XL developed code to analyze screening or characterization data. TT, AJM, MAL, XL and AM benchmarked GEVIs *in vitro*. FSP, AJM, MAL, and SY prepared figures and wrote the manuscript. **Bourdieu lab.** VV, JB, and LB coordinated the project and wrote the manuscript. AA, DGS and JB generated AAVs. AL and VV designed optimized ULoVE patterns. BM implemented diagonal scanning. VV and DGS performed mouse surgeries, conducted experiments, analyzed data, and prepared figures. **Ji lab.** NJ supervised the project. JZ prepared samples, acquired and analyzed data, and made figures. RGN performed mouse surgeries. HY analyzed data. All authors contributed to the writing of the manuscript. **Reimer lab.** JR supervised the project. MG, NH, and RK collected the optical data. RGL, MG, NH, YYS, and MH performed the analyses and generated the figures. JR, MG, and NH wrote the manuscript. **Zhong lab.** WZ supervised the project. MP conducted and analyzed the experiments.

