## Supplementary Information for "A versatile, positive-going voltage indicator that enables accessible two-photon recordings in vivo"

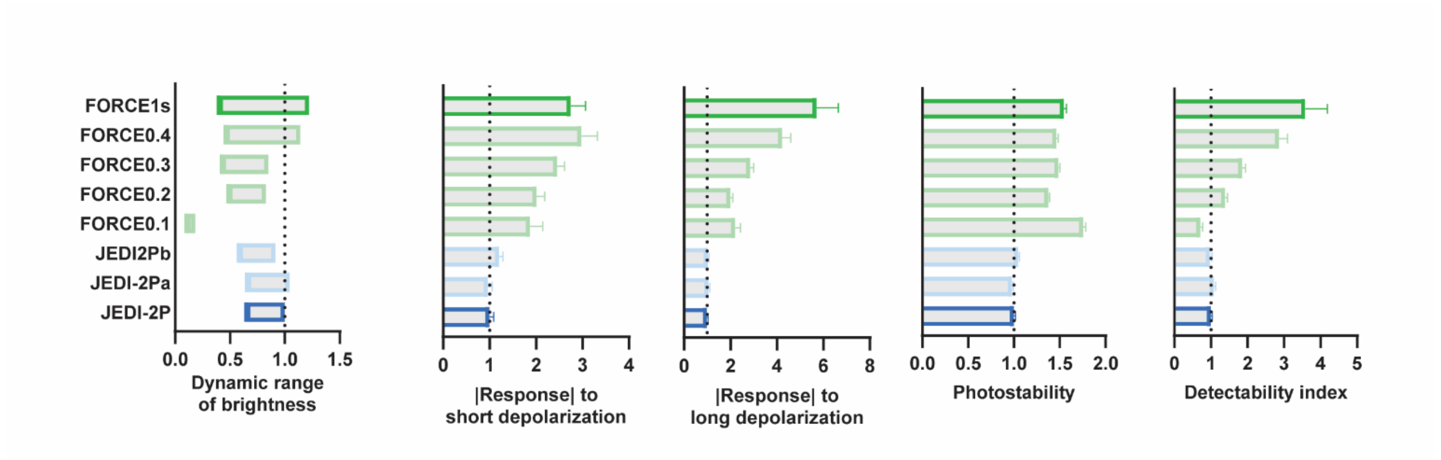


**Figure S1.1. Detectability gains during FORCE1s directed evolution**.

Performance of FORCE1s and intermediate variants across the 2P screening metrics, shown relative to JEDI-2P. The dynamic range of brightness reflects the span between the dimmest and brightest GEVI signals observed during the long electric-field stimulation protocol (5 × 10-ms pulses of 30 V). |Response| is the peak $|\Delta F/F_0|$ achieved during short (1 ms, 60 V) and long electric-field stimulations. Photostability is the area-under-the-curve of baseline GEVI brightness across the 9-second screening protocol. The detectability index is calculated as $|\Delta F/F_0|_{long} * \sqrt{B}$, where $|\Delta F/F_0|_{long}$ is the response to long electric-field stimulations and B is the baseline brightness, calculated by normalizing GEVI baseline fluorescence ($F_0$) with fluorescence values from the covalently attached mBeRFP. By integrating response amplitude and brightness, the detectability index serves as a composite metric for ranking indicators during screening, offering a practical estimate of signal-to-noise ratio in voltage transient detection. n = 6 independent transfections per variant. Error bars: 95% CI.

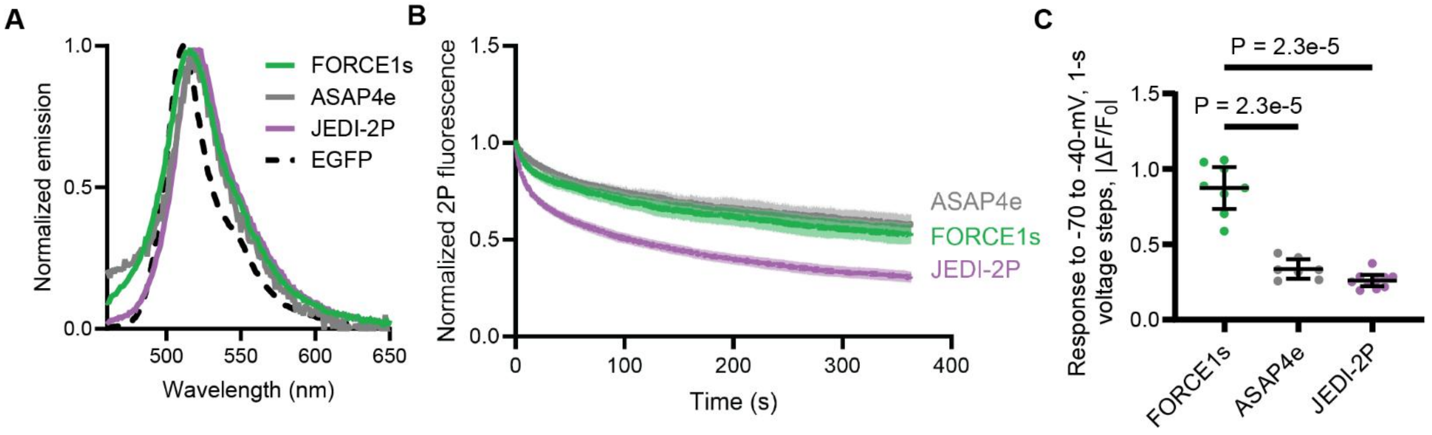


**Figure S1.2. Characterization of FORCE1s emission spectrum, photostability, and steady-state responses.**
(**A**) Emission spectra. Mean emission peaks ± 95% CI, ranked from blue to red, were: 511 ± 1 (EGFP), 516 ± 5 (FORCE1s), 518 ± 4 (ASAP4e), and 519 ± 3 nm (JEDI-2P). Constructs were expressed in HEK293A cells, and excited at 430/10 nm (mean/bandwidth). Spectra were normalized to their respective peaks and averaged from n = 6 (FORCE1s) or 3 (other constructs) independent transfections. To ensure consistent expression and analysis conditions, EGFP was targeted to the plasma membrane via a C-terminal prenylation sequence (CAAX). (**B**) FORCE1s exhibited greater photostability than JEDI-2P, and comparable photostability to ASAP4e (see Sup. Stats). GEVIs were imaged with 100 mW power at the sample plane 940-nm light for 6 min using 2P resonance-scanning over a 40 x 635 µm (32 rows x 512 columns) FOV. Fluorescence was normalized to the first frame. Dark traces: mean from n = 12 independent transfections per GEVI. Shaded regions: 95% CI. (**C**) Quantification of (Fig. 1E) for -70 to -40 mV voltage steps. Black lines: mean. Error bars: 95% CI. P-value corresponds to Dunnett's T3 multiple comparisons test between FORCE1s and each of the other two GEVIs, conducted after a significant Welch ANOVA test.

| @ $-83.1 \pm 0.5$ mV | FORCE1s | JEDI-2P |
|---|---|---|
| $\tau$ fast (ns) | $1.3 \pm 0.1$ | $1.7 \pm 0.2$ |
| $\tau$ slow (ns) | $3.4 \pm 0.0$ | $3.4 \pm 0.1$ |
| % slow | $88 \pm 1$ | $79 \pm 5$ |

| @ $-10.5 \pm 0.1$ mV | | |
|---|---|---|
| $\tau$ fast (ns) | $1.3 \pm 0.2$ | $1.5 \pm 0.1$ |
| $\tau$ slow (ns) | $3.5 \pm 0.0$ | $3.3 \pm 0.0$ |
| % slow | $89 \pm 1$ | $80 \pm 2$ |

**Table S1.3. Indicator fluorescence lifetime at 22°C.**
The resting membrane potential was modulated by changing the external potassium concentration, from 5 mM (*top*, hyperpolarized) to 100 mM (*bottom*, depolarized). Measurements were acquired. Lifetime values are the mean ± 95% CI. n = 95 (FORCE1s) and 89 (JEDI-2P) HEK Kir2.1 cells. Membrane potential values are mean ± SEM, n=4 HEK Kir2.1 cells.

| -70 to +30 mV | FORCE1s | ASAP4e | JEDI-2P |
|---|---|---|---|
| $\tau$ **fast (ms)** | 2.8 ± 0.1 | 2.6 ± 0.8 | 0.49 ± 0.07 |
| $\tau$ **slow (ms)** | 16 ± 2 | 11 ± 2 | 2.4 ± 0.8 |
| **% fast** | 93 ± 2 | 34 ± 10 | 78 ± 9 |

| +30 to -70 mV | FORCE1s | ASAP4e | JEDI-2P |
|---|---|---|---|
| $\tau$ **fast (ms)** | 2.4 ± 0.6 | 6.7 ± 0.5 | 1.5 ± 0.1 |
| $\tau$ **slow (ms)** | 9.1 ± 1.0 | n/a[a] | n/a[a] |
| **% fast** | 47 ± 8 | n/a[a] | n/a[a] |

**Table S1.4. Indicator kinetics at 33°C under widefield 1P illumination**.
The voltage of HEK293A cells was modulated under whole-cell voltage clamp. Values are the mean ± 95% CI. n = 14 (FORCE1s), 17 (ASAP4e), and 11 (JEDI-2P) cells.

[a] These kinetics were best fit by a monoexponential function.

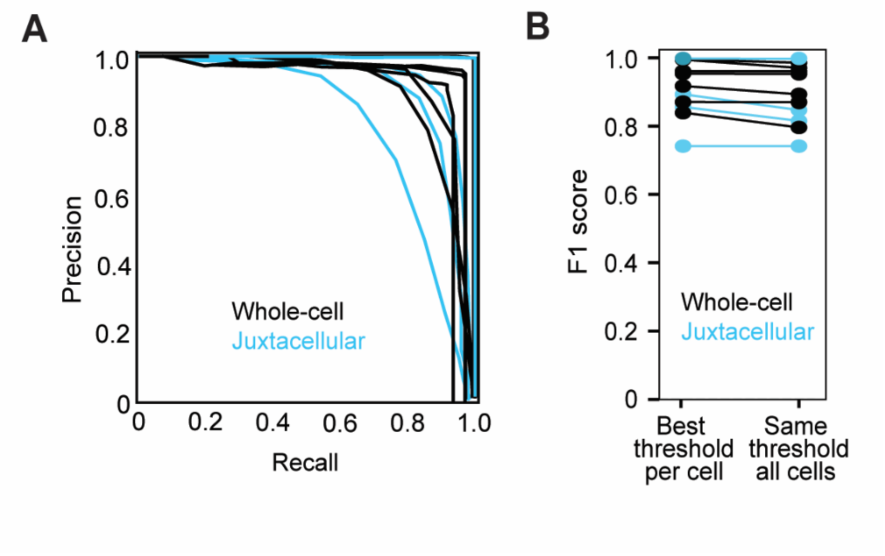


**Figure S2.1. Precision-recall analysis combining whole-cell and juxtacellular electrophysiological recordings.** Simultaneous imaging was conducted using 440 Hz resonant scanning. (**A**) Precision-recall curves. n = 11 neurons, combining 7 whole-cell (black) and 4 juxtacellular (blue) patch-clamp measurements. (**B**) F1 scores for the 11 cells from panel A. Best threshold per cell: 0.91±0.02 (mean ±SEM). Same threshold all cells: 0.89±0.03 (mean ±SEM).

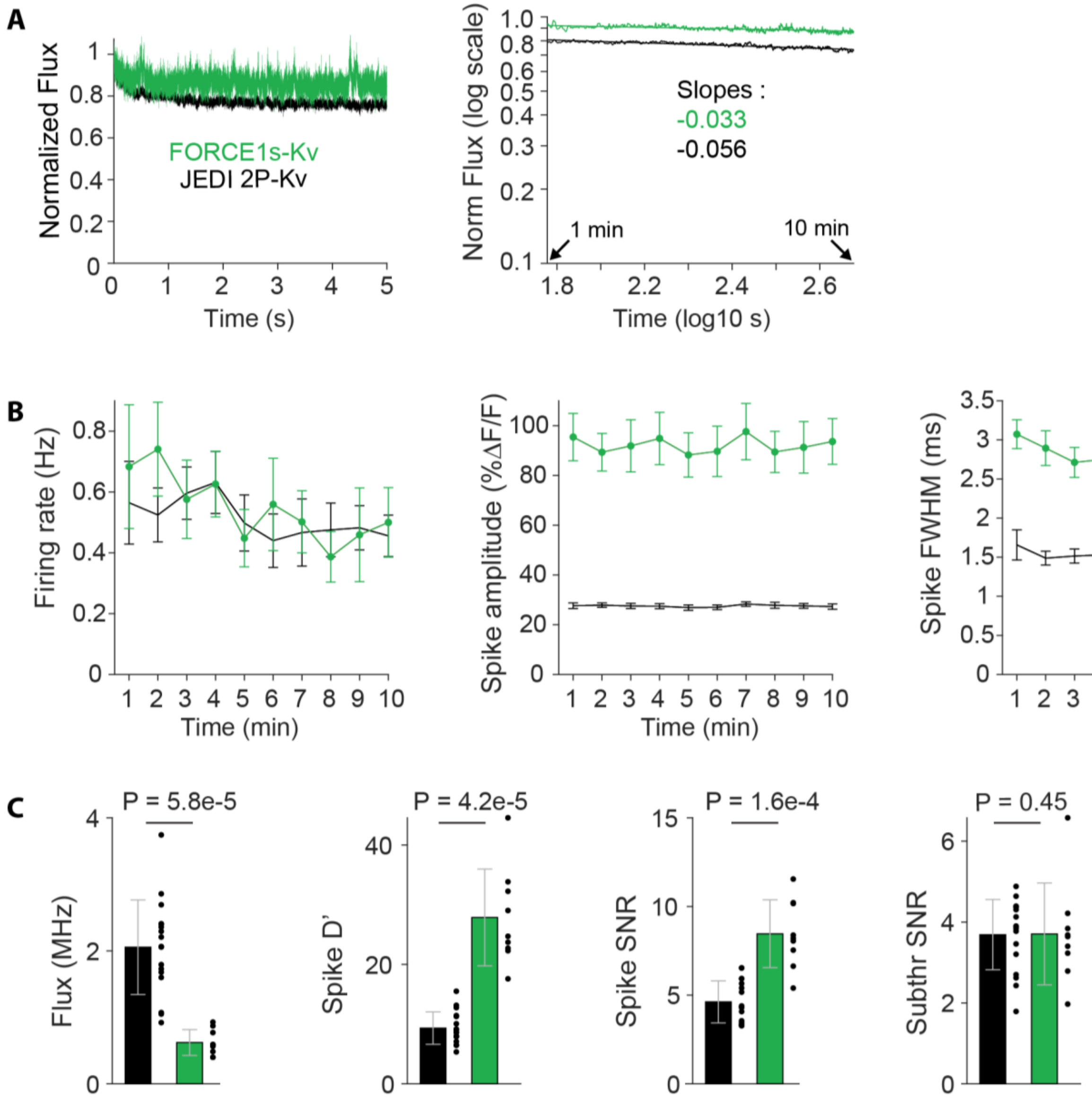


**Figure S3.1. Additional quantitative comparisons of the performance of JEDI2P-Kv and FORCE1s-Kv.**
(**A**) Photostability quantification. FORCE1s-Kv is less prone to photobleaching as evidenced by a lower change in average photon flux during the initial fast bleaching phase (*left*) and by a lower power-law slope in the remaining fluorescence trace (*right*). n =17 (JEDI2P-Kv), 9 (FORCE1s-Kv) neurons. (**B**) Quantification of spiking properties over the duration of recordings. Datapoints: mean values. Error bars: SEM. n =17 (JEDI2P-Kv), 9 (FORCE1s-Kv). Firing rate decreased by 0.01472 (JEDI-2P-Kv) and 0.02925 (FORCE1s-Kv) Hz per min of recordings. (**C**) Quantitative comparisons. Error bars: SD. n =17 (JEDI2P-Kv), 9 (FORCE1s-Kv) neurons. P-values from two-sample Wilcoxon rank sum test.

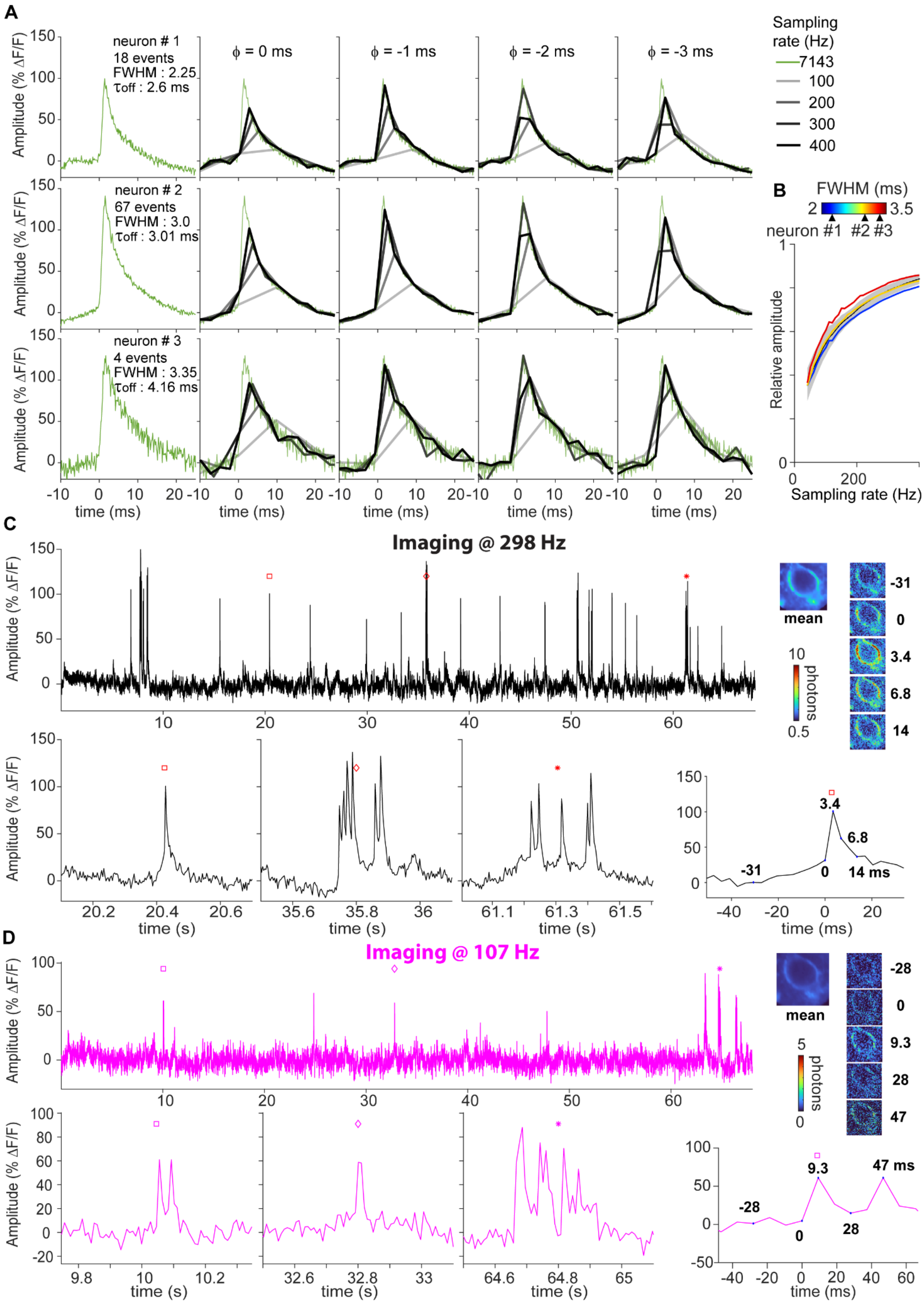

A
neuron # 1
18 events
FWHM : 2.25
τoff : 2.6 ms
neuron # 2
67 events
FWHM : 3.0
τoff : 3.01 ms
neuron # 3
4 events
FWHM : 3.35
τoff : 4.16 ms
ϕ = 0 ms
ϕ = -1 ms
ϕ = -2 ms
ϕ = -3 ms
Amplitude (% ΔF/F)
time (ms)
Sampling rate (Hz)
7143
100
200
300
400
B
FWHM (ms)
2
3.5
neuron #1 #2 #3
Relative amplitude
Sampling rate (Hz)
C
Imaging @ 298 Hz
Amplitude (% ΔF/F)
mean
photons
-31
0
3.4
6.8
14
time (s)
time (ms)
14 ms
D
Imaging @ 107 Hz
Amplitude (% ΔF/F)
mean
photons
-28
0
9.3
28
47
time (s)
time (ms)
47 ms

**Figure S3.2. Sampling-rate dependence of FORCE1s-Kv spike amplitudes using ULoVE and diagonally-scanned imaging.**

(**A**) In silico evaluation of sampling frequency effects of spike waveform recordings with ULoVE. Dwell time was held constant; sampling rates below 7.1 kHz represent downsampling. The leftmost column shows average spike waveforms from three representative neurons with mean spike widths of 2.25, 3.00, and 3.35 ms (top to bottom). Subsequent columns display predicted waveforms at different sampling frequencies (colored traces) and corresponding phase delays relative to spike onset (Φ). FWHM, Full Width at Half Maximum. (**B**) Plots of relative spike amplitude versus sampling rate for the mean (black) of the 9 FORCE1s neurons ± SD (grey), including those shown in panel A and color-coded by their FWHM (#1, #2, #3), recorded at 7.1 kHz using ULoVE. (**C-D**). A representative neuron recorded at (**C**) 298 Hz and (**D**) 107 Hz using diagonally-scanned imaging, illustrating how insufficient sampling can substantially reduce the peak response amplitude to spikes. Zoom-in epochs below show isolated action potentials and spike bursts at the times indicated by symbols in the upper trace. *Top right*: mean image of the neuron alongside individual frames during a spike. *Bottom right*: zoom-in epoch of the same spike shown in the top right time-lapse.

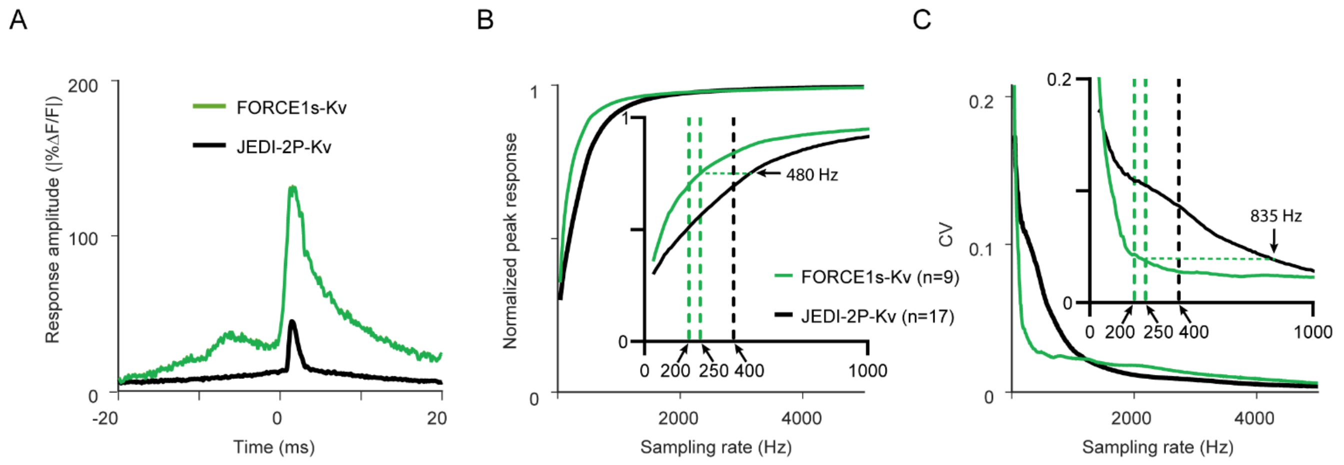


**Figure S3.3. Sampling-rate dependence of spike amplitude and variability in ULoVE: FORCE1s-Kv versus JEDI-2P-Kv.** (**A**) Average spike responses from individual neuronal recordings shown in Fig. 3A (n = 126 and 295 spikes for FORCE1s-Kv and JEDI-2P-Kv, respectively). (**B**) Relative peak spike amplitude (mean over 20 phases) of the 9 FORCE1s-Kv and the 17 JEDI-2P-Kv expressing neurons plotted against sampling rate (down-sampled from their 1MHz up-sampled average spike waveforms). Inset: a zoom-in from 0 to 1000 Hz. The equivalent normalized peak response of JEDI-2P to FORCE1s when sampled at 250 Hz is indicated with a horizontal dashed line. **(C)** Coefficient of variation (CV), corresponding to the SD over the mean, plotted against sampling rate. The equivalent CV of JEDI-2P to FORCE1s when sampled at 250 Hz is indicated with a horizontal dashed line.

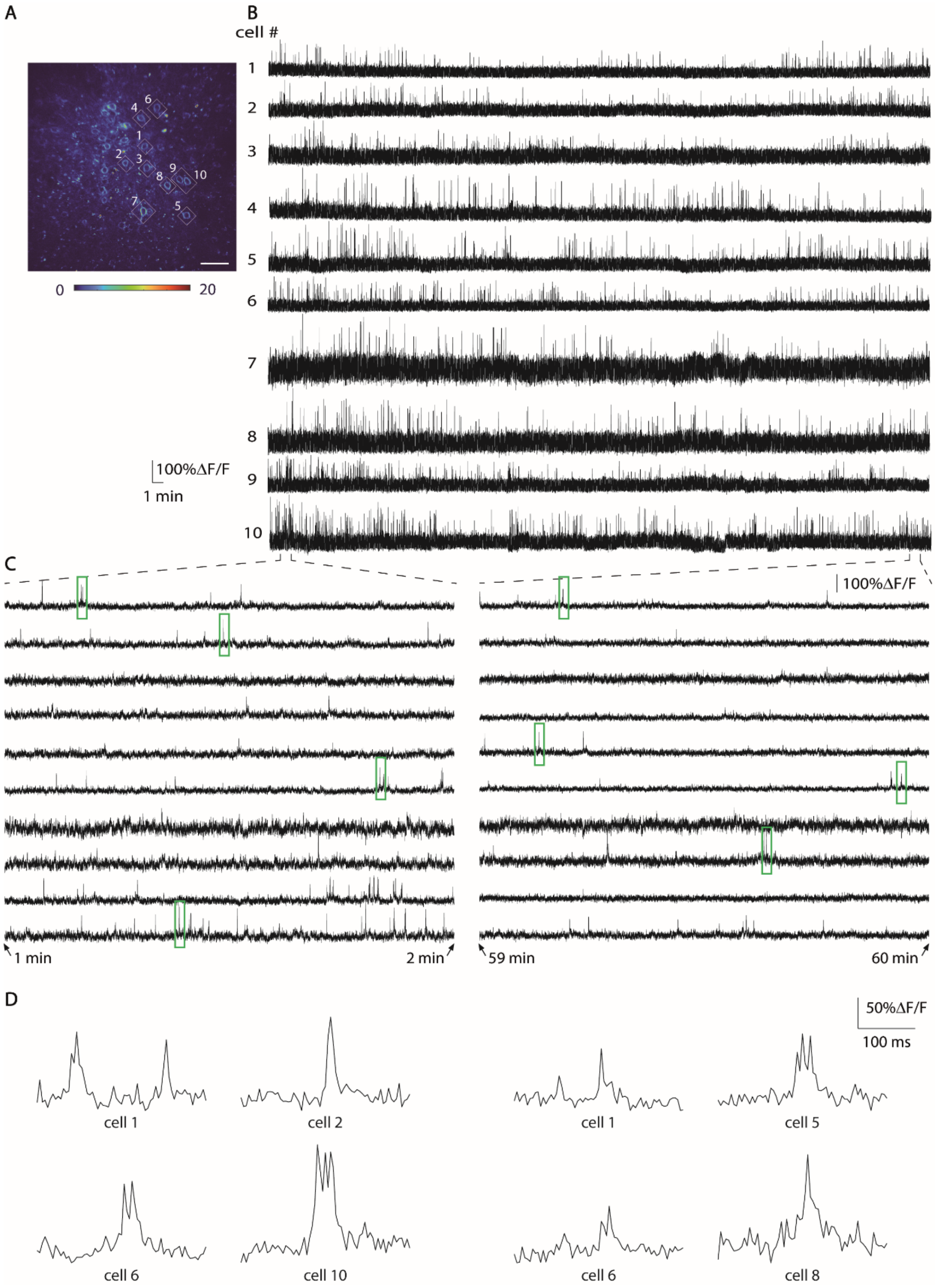


**Figure S3.4. Hour-long, multi-cell recordings with diagonal scanning – Recording 1.**
(**A**) FORCE1s-Kv-expressing neurons in layer 2/3 of V1 visual cortex. White lozenges mark scanned regions. Image is a mean-intensity projection from a 20-frame time series. Colormap indicates minimum and maximum gray values. Resolution: 0.27 µm/pixel. Scale bar: 50 µm. (**B**) 61-min recordings from the neurons highlighted in A, acquired at 217 Hz with a dwell time of 0.25 µs/pixel and resolution of 0.91 µm/pixel. (**C**) One-minute zoomed segments from traces in B. (**D**) 300-ms zoomed epochs corresponding to green rectangles in panel C.

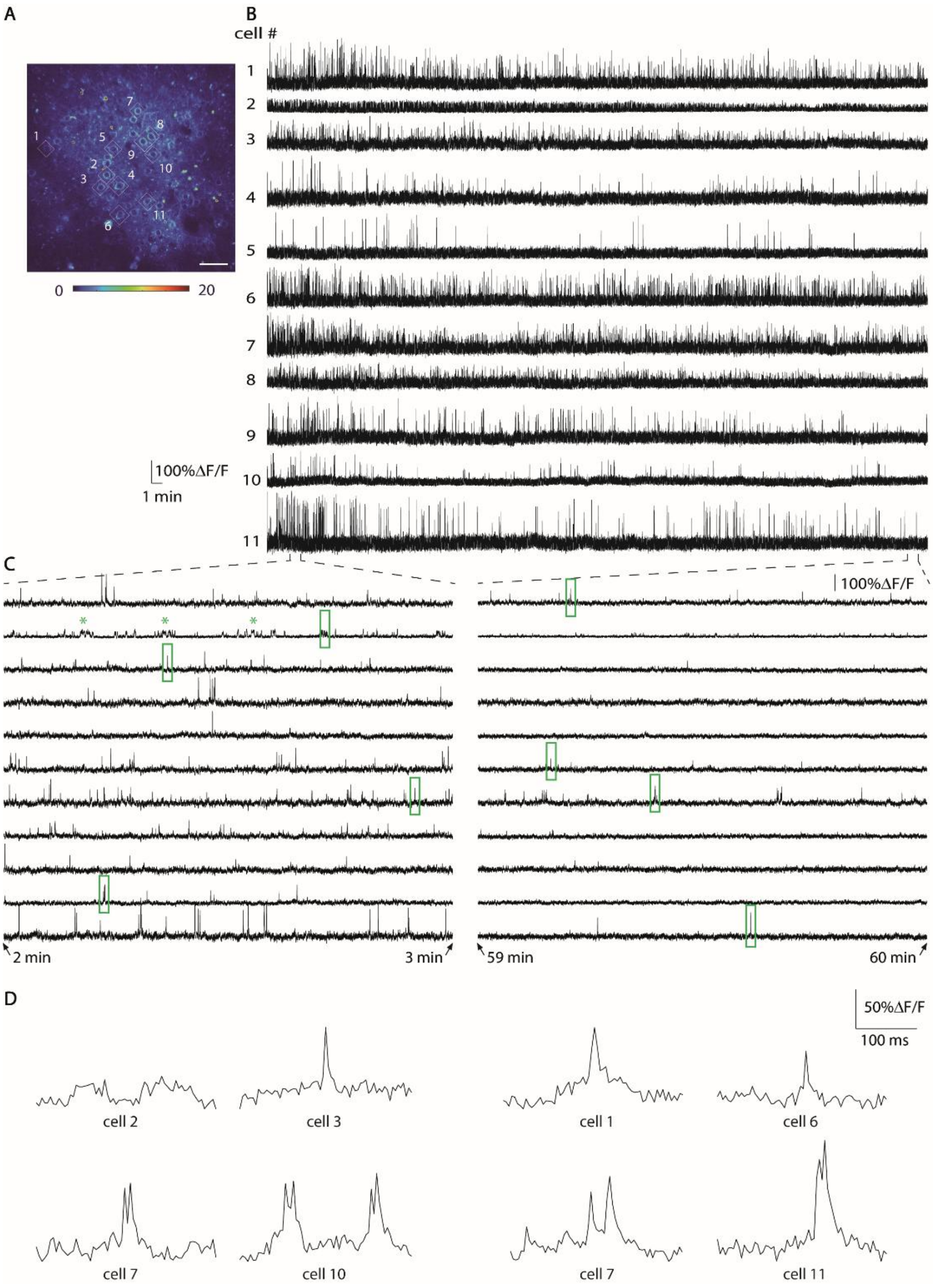


**Figure S3.5. Hour-long, multi-cell recordings with diagonal scanning – Recording 2.**

(**A**) FORCE1s-Kv–expressing neurons in layer 2/3 of V1 visual cortex. White lozenges mark scanned regions. Image is a mean-intensity projection from a 20-frame time series. Colormap indicates minimum and maximum gray values. Resolution: 0.27 µm/pixel. Scale bar: 50 µm. (**B**) 61-min recordings from the neurons highlighted in A, acquired at 214 Hz with a dwell time of 0.25 µs/pixel and resolution of 0.91 µm/pixel. (**C**) One-minute zoomed segments from traces in B. Note that neuron #2 was found to have a pattern of long depolarizing plateaus (asterisks) and an absence of spiking activity of expected amplitudes. This phenotype occurred in 1 out of 49 recorded cells. (**D**) 300-ms zoomed epochs corresponding to green rectangles in panel C.

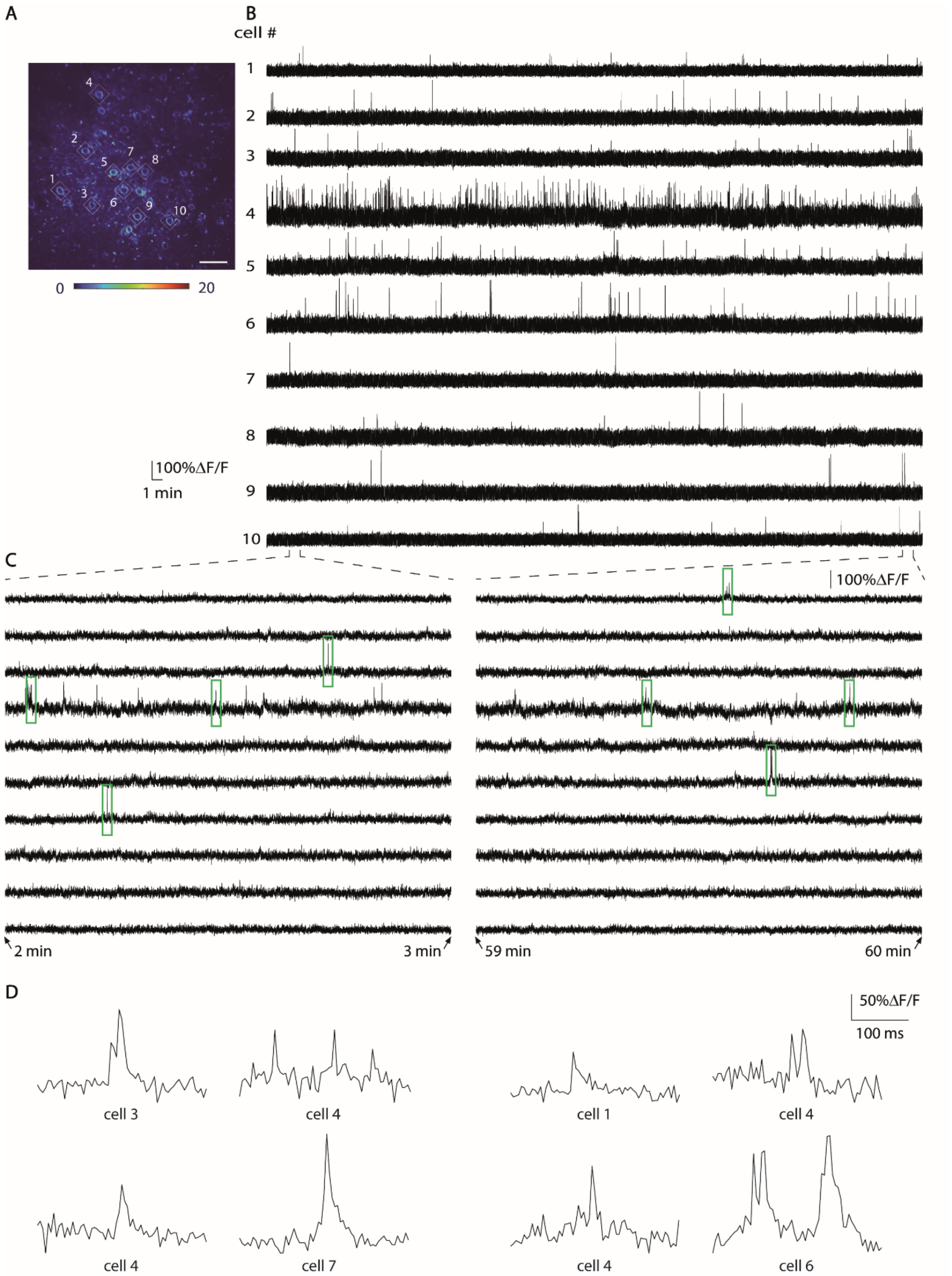


**Figure S3.6. Hour-long, multi-cell recordings with diagonal scanning – Recording 3.**
(**A**) FORCE1s-Kv–expressing neurons in layer 2/3 of V1 visual cortex. White lozenges mark scanned regions. Image is a mean-intensity projection from a 20-frame time series. Colormap indicates minimum and maximum gray values. Resolution: 0.27 μm/pixel. Scale bar: 50 μm. (**B**) 61-min recordings from the neurons highlighted in A, acquired at 210 Hz with a dwell time of 0.25 μs/pixel and resolution of 0.91 μm/pixel. (**C**) One-minute zoomed segments from traces in B. (**D**) 300-ms zoomed epochs corresponding to green rectangles in panel C.

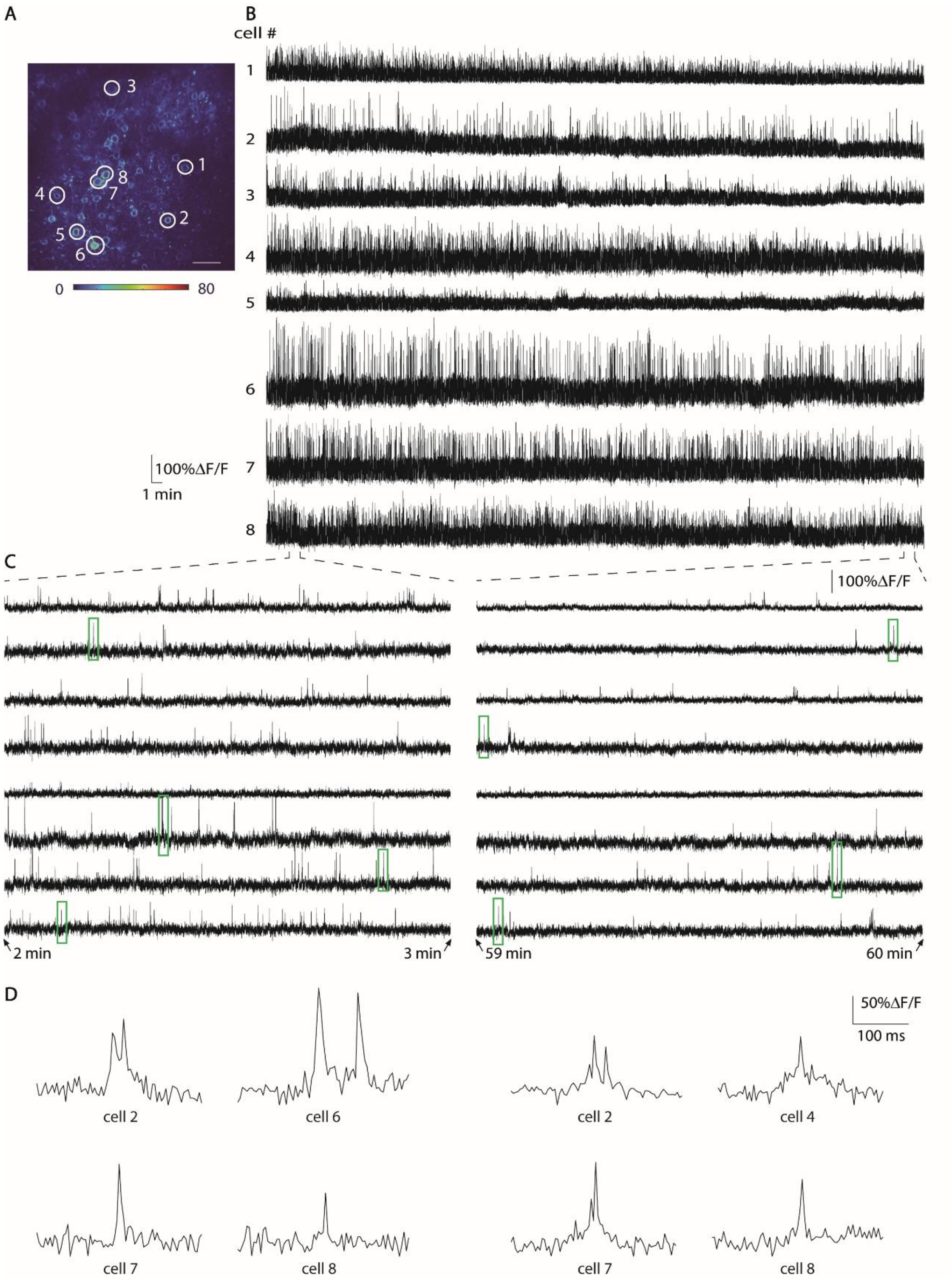


**Figure S3.7. Hour-long, multi-cell recordings with diagonal scanning – Recording 4.**
(**A**) FORCE1s-Kv–expressing neurons in layer 2/3 of V1 visual cortex. White outlines mark scanned regions. Image is a z-stack of 10 frames around the imaged plane at 0.18 µm/ pixel resolution. Colormap indicates minimum and maximum gray values. Resolution: 0.18 µm/pixel. Scale bar: 50 µm. (**B**) 61-min recordings from the neurons highlighted in A, acquired at 250 Hz with a dwell time of 0.20 µs/pixel and resolution of 0.91 µm/pixel. (**C**) One-minute zoomed segments from traces in B. (**D**) 300-ms zoomed epochs corresponding to green rectangles in panel C.

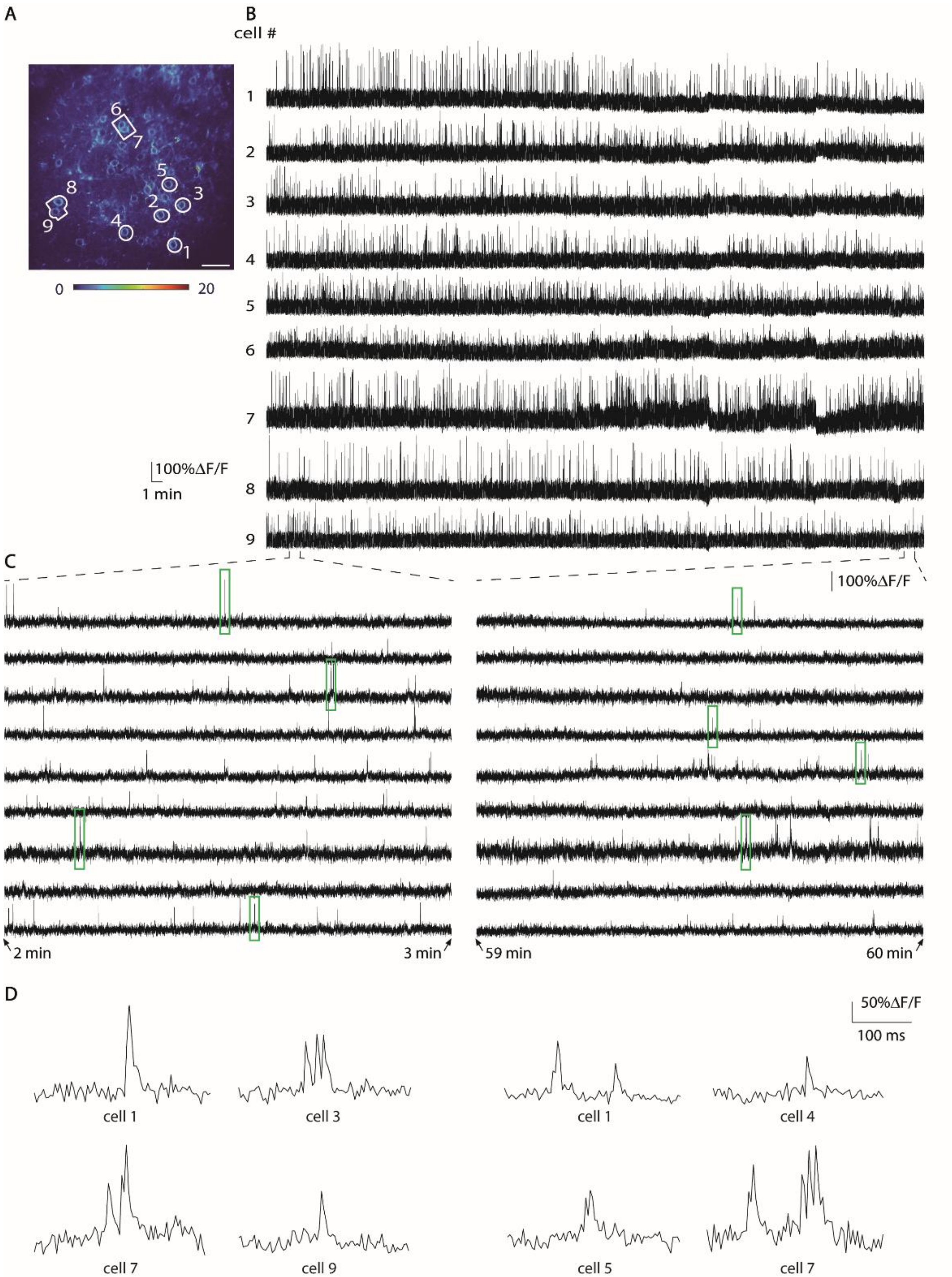


**Figure S3.8. Hour-long, multi-cell recordings with diagonal scanning – Recording 5.**

(**A**) FORCE1s-Kv–expressing neurons in layer 2/3 of V1 visual cortex. White outlines mark scanned neurons. Image is a mean-intensity projection from a 7-frame time series. Colormap indicates minimum and maximum gray values. Resolution: 0.18 µm/pixel. Scale bar: 50 µm. (**B**) 61-min recordings from the neurons highlighted in A, acquired at 257 Hz with a dwell time of 0.275 µs/pixel and resolution of 1.0 µm/pixel. (**C**) One-minute zoomed segments from traces in B. (**D**) 300-ms zoomed epochs corresponding to green rectangles in panel C.

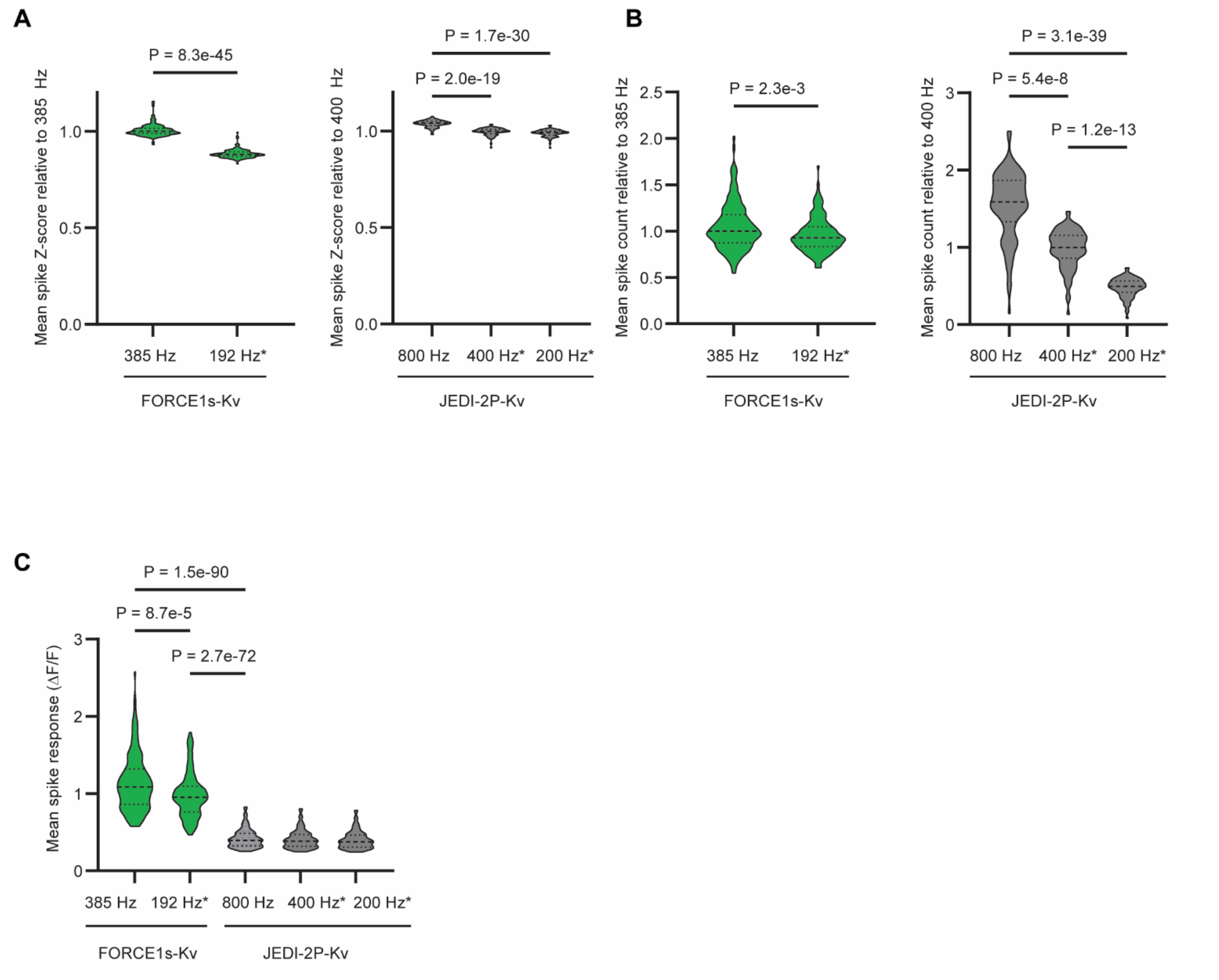


**Figure S4.1 Sampling-rate dependence of spike detection with FACED2.0 imaging of FORCE1s-Kv.**
All plots were truncated to their maximum and minimum values, and the asterisk (*) indicates downsampled datasets. (**A**) Violin plots of relative spike Z-scores of FORCE1s-Kv (left, green) and JEDI-2P-Kv (right, gray) recordings compared with the same traces downsampled 2x or 4x. The JEDI-2P-Kv recordings are from our previous study. n = 134 (FORCE1s-Kv) and 84 (JEDI-2P-Kv) neurons from one FOV per construct. Dashed lines are the median (thick lines) and quartiles (thin lines). P-values are from Mann-Whitney (left) and Kruskal-Wallis (right) tests. (**B**) Violin plots of relative spike counts from the same recordings analyzed in panel A. P-values are from a lognormal Welsh's t-test (left) or a Kruskal-Wallis test (right). (**C**) Violin plots of average spike responses (ΔF/F) from the same recordings analyzed in panel A. P-values are from Šídák's multiple comparisons tests.

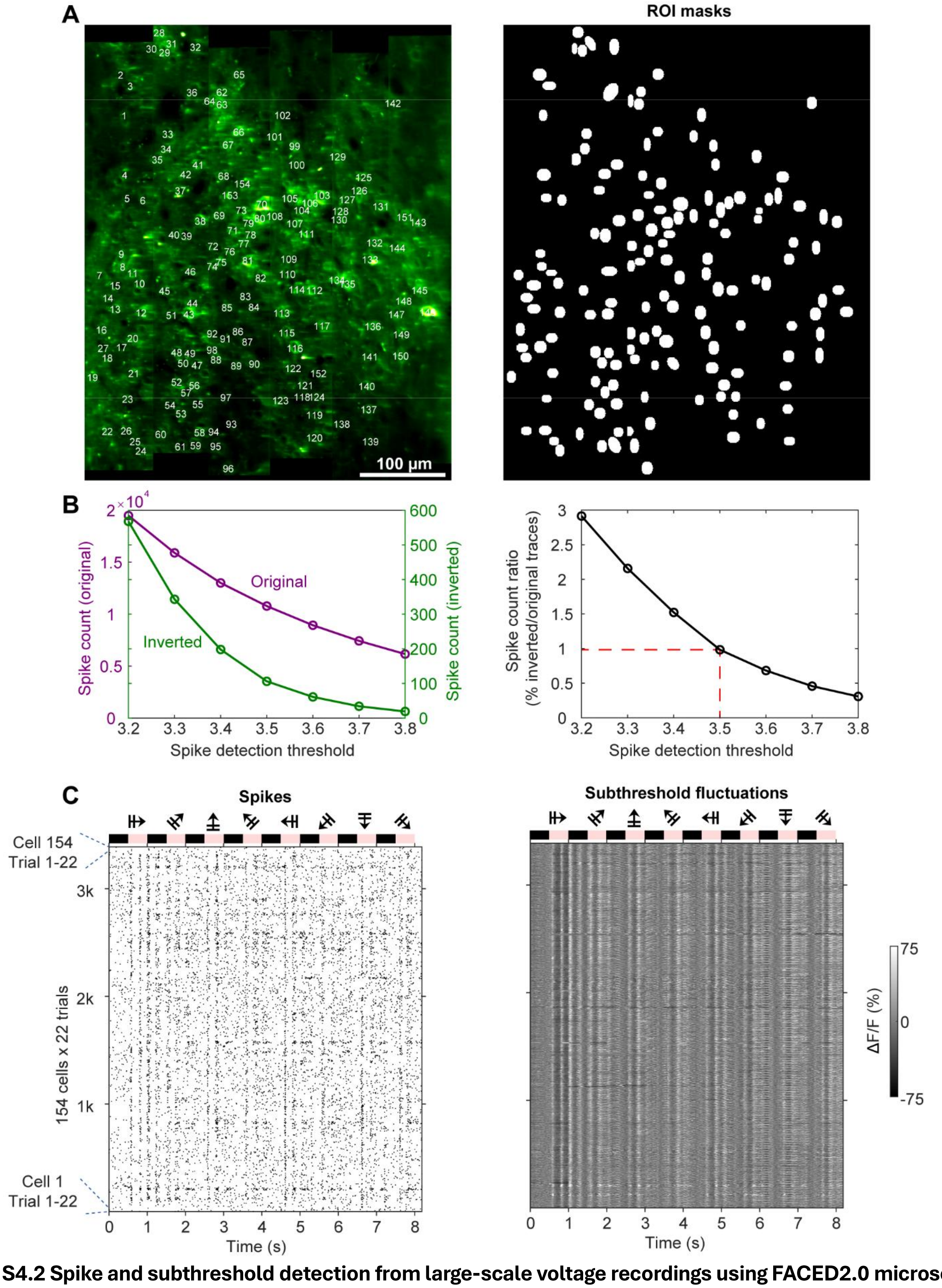


**Figure S4.2 Spike and subthreshold detection from large-scale voltage recordings using FACED2.0 microscopy.** (**A**) Regions of interest (ROI) indices (left) and ROI masks (right) from the neurons shown in Fig. 4B. (**B**) Left. Detected spike counts as a function of the detection threshold, expressed as the Z-score of the high-pass-filtered trace (purple). False positives were estimated by repeating the analysis on inverted traces (green). Right: Ratio of spikes detected in inverted versus original traces. The red dashed lines show the chosen threshold of 3.5 giving a false-positive rate of 1%. (**C**) Spike raster plot (left) and subthreshold fluctuations (right) from all cells and trials, grouped by cells. Pink blocks above the plots indicate periods of grating stimulus presentation; symbols above denote stimulus orientation.
